# Assessing the role of mitonuclear interactions on mitochondrial function and organismal fitness in natural *Drosophila* populations

**DOI:** 10.1101/2023.09.25.559268

**Authors:** S Bettinazzi, J Liang, E Rodriguez, M Bonneau, R Holt, B Whitehead, D.K Dowling, N Lane, MF Camus

## Abstract

Mitochondrial metabolism is regulated by a series of enzyme complexes, whose function depends on effective interactions between proteins and RNA encoded by the mitochondrial and nuclear genomes. These epistatic interactions are in turn highly sensitive to the environment. Many studies have found that mitochondrial haplotype frequencies of various taxa associate with latitude or altitude, leading to the hypothesis that mitochondrial genomes may respond to thermal selection and contribute to local adaptation. We used a *Drosophila melanogaster* panel comprising native (coadapted) populations from the extremes of the Australian east-coast cline, and generated mitonuclear cybrid populations. Our results indicate a strong phenotypic impact of mitonuclear interactions in cybrid lines, involving an apparent trade-off between aerobic capacity and key fitness aspects such as reproduction, growth, and survival. Overall, our study shows that naturally-occurring mitonuclear disruptions can have a meaningful impact on phenotypes, potentially influencing future ecological adaptation and population persistence.

## Introduction

Metabolism lies at the core of life-history theory (Burger *et al*. 2019). To thrive, organisms must adapt and exploit the resources offered by the environment. Mitochondria are key for metabolic adaptation, as they are central hubs for both energy transduction and intermediary metabolism in eukaryotes — a major determinant of all aspects of fitness including growth, development, and reproductive success (Lane 2009). Despite its key role, oxidative phosphorylation is uniquely vulnerable to disruption, as it depends on components encoded by two different genomes, mitochondrial (mt) and nuclear (n), which must interact harmoniously with each other to preserve bioenergetic efficiency (Blier *et al*. 2001; Dowling *et al*. 2008; Lane 2009, 2011; Wolff *et al*. 2014). Whilst coevolution is vital, mitonuclear interactions are perpetually stressed by their very different evolutionary dynamics. As the mitochondrial genome is typically uniparentally inherited and mutates at a faster rate than the nuclear genome across many species, it is thus predicted that the recombining nuclear genome will be able to respond adaptively to its fast-changing mitochondrial counterpart (Barreto *et al*. 2018; Healy & Burton 2020; Hill 2020). This process has been shown to result in tight mitonuclear coevolution in some taxa, which has the potential to lead to rapid divergence between populations in their mitonuclear genotypes (Morales *et al*. 2018; Wang *et al*. 2021; Burton 2022; Biot-Pelletier *et al*. 2023). This process poses a challenge in the event of admixture between disjunct populations with unique trajectories of mitonuclear coevolution, as they may generate genomic incompatibilities.

Environmental changes are occurring at an unprecedented pace, with large consequences on species persistence and distributions (Outhwaite *et al*. 2022). Many species are predicted to have shrinking or altered distributions, requiring adaptation to new environments or migration to more favourable habitats. This process could lead to divergent populations having to re-unite and share a common niche. Although new mitonuclear variants may foster evolutionary innovations and be favoured under certain conditions (e.g. providing better adaptation to the environment), admixture may also unmask mitonuclear incompatibilities and manifest as decrease in life-history trait expression. There are examples of detrimental mitonuclear interactions impacting all aspects of individual fitness in yeast (Lee *et al*. 2008; Biot-Pelletier *et al*. 2023), invertebrates (Sackton *et al*. 2003; Ellison & Burton 2006; Ellison *et al*. 2008; Meiklejohn *et al*. 2013; Zhang *et al*. 2017; Camus *et al*. 2020b; Rank *et al*. 2020) and vertebrates (Barrientos *et al*. 1998; Dey *et al*. 2000; Deremiens *et al*. 2015; Ma *et al*. 2016; Chapdelaine *et al*. 2020). However, most of the studies to date have focused on laboratory crosses between inbred populations, or interspecific crosses between geographically distant species, making the results difficult to translate to real-world population ecology.

Here we examine the contribution of mitonuclear genotypes to locally adaptive phenotypes, combining mitochondrial physiology with life-history trait expression. Using *Drosophila* populations sampled from low and high latitude localities of the Australian east coast (locally adapted to subtropical and temperate environments), we created a full-factorial mitonuclear population panel; exploiting genetic variation naturally present to test the effect of natural admixture events along the cline. Following a mitonuclear coadaptation hypothesis, we predicted that native fly lines would have higher fitness than cybrid lines with a higher proportion of disrupted mitonuclear alleles. The physiological impact of mitonuclear breakdown was assessed for both mitochondrial bioenergetics and metabolically important life-history traits. Our results demonstrate that even mild mitonuclear divergence, of the kind resulting from natural admixture events, can strongly impact the organismal phenotype. Specifically, cybrid lines exhibited an apparent trade-off between aerobic capacity and key fitness aspects, characterized by increased OXPHOS activity, lower ROS production and higher motility, but decreased reproductive success, growth rate and survival compared to parental populations. Overall, our study supports the idea that intergenomic interactions can be a strong determinant of individual fitness, adaptive capacity, and population persistence.

## Materials and Methods

Materials and methods are described briefly. Further detailed information can be found in *Supporting material*.

### Mitonuclear panel establishment and maintenance

Two replicated *Drosophila melanogaster* populations were sourced from the Australian east coast cline in early 2021. These populations were from Townsville ‘T’ (latitude: -19.26, longitude 146.81) and Melbourne ‘M’ (latitude: -37.77, longitude: 144.99). Following the creation of these massbred populations, a full factorial mitonuclear panel was generated. The panel followed a two-letter nomenclature, indicating the mitochondrial genome first and the nuclear background second (lower and uppercase letter, respectively). It included two parental populations with naturally coevolved mitonuclear genotypes, named after the nDNA sampling area (i.e. ‘tT’ - Townsville and ‘mM’ - Melbourne), and two derived cybrid lines (‘mT’ and ‘tM’), where the mitochondrial and the nuclear background were reciprocally swapped, using a well-established balancer chromosome crossing scheme (figure s1) (Zhu *et al*. 2014; Camus *et al*. 2017b). Given the turnover of *Drosophila* is roughly a fortnight, replicates of all four populations were setup in such a way that we would have flies emerging on a weekly basis (this term is referred to as “*pop*” in statistical analyses).

Experimental lines were maintained at standard laboratory conditions (25°C, 50% RH, 1:1 protein:carbohydrate P:C diet, 12:12 light:dark day cycle). Mitochondrial DNA congruence was routinely checked for by means of PCR, whereas nuclear genetic variance was preserved by regular backcrossing to the nuclear-correspondent massbred native lines. Prior to each experiment, flies were reared in density-controlled conditions (20 eggs per vial), sorted by sex 48-hours post eclosion (excluding reproductive performance), let to acclimate in new food vials and finally assayed at 4-7 days of age. Reproductive performance assays used the same rearing scheme for focal flies, however experimental flies were collected as virgin (within 2-5 hours post eclosion).

### Mitochondrial physiology

Mitochondrial bioenergetics were characterized at 25°C on fly permeabilized tissue using dedicated Oxygraph-2k-FluoRespirometers (Oroboros Instruments, Innsbruck, Austria), following existing protocols with minor modifications (Bettinazzi *et al*. 2019; Gnaiger 2020; Rodríguez *et al*. 2021; Rodríguez *et al*. 2023). Following a specific SUIT protocol (see supporting information), we assessed mitochondrial respiration sustained by different combination of respiratory complexes and in different respiratory states. This included the activity of complex I (CI), proline dehydrogenase (ProDH), complex II (CII), glycerophosphate dehydrogenase (GpDH) and complex IV (CIV), as well as respiratory states such as Leak (non-phosphorylating resting state - state 4 or 2’), OXPHOS (coupled respiration – state 3) and ETS (uncoupled respiration, state 3u). H2O2 fluxes were evaluated in parallel with respiratory rates, and parameters named accordingly. Respirometry data were expressed as O2 fluxes normalized for tissue mass (pmol O2 · s^−1^ · mg^−1^), and as flux control ratios (FCR), normalized for maximum coupled respiration (CI+ProDH+CII+GpDHP) (Gnaiger 2020). Change in respiration following the addition of specific substrates or inhibitors was expressed by means of flux control factors (FCF). Reactive oxygen species rates were expressed as H2O2 fluxes normalized for tissue mass (pmol H2O2 · s^−1^ · mg^−1^), or as ratios, in function of the step-specific oxygen consumption (H2O2·O ^−1^).

### mtDNA copy number

Genome abundance was determined fluorometrically on a Mastercycler® RealPlex thermocycler (Eppendorf, DE), using the KAPA SYBR® FAST qPCR Master Mix Kit (KAPABIOSYSTEMS) and two complementary sets of primers, respectively amplifying *cox1* (mitochondrial) and *rosy* (nuclear) genes. For both genes, the cycle thresholds (CT) were measured in duplicates and the mtDNA copy number relatively to the nuclear genome determined by the formula (2^-ΔCT^)·2, with ΔCT referring to the difference between the mitochondrial and the nuclear gene mean CT (Ballard *et al*. 2007b).

### Locomotor activity

Fly locomotor activity was recorded for 48 hours using dedicated Drosophila Activity Monitors (DAM2, Trikinetics), and calculated as the number of counts (infrared beam breaks) per minute (Anderson *et al*. 2022). Activity was then condensed in 30 minutes activity, and further in timeframe-specific activity (i.e. dawn, day, dusk and night).

### Reproductive performance

Reproductive fitness was investigated in both female and male individuals, the latter in both a non-competitive and competitive environment. All adults were collected as virgins (within 5 hours post-eclosion), reared and acclimated at the same standard laboratory conditions (25°C, 50% RH, 1:1-P:C diet, 12:12 light:dark cycle) and of the same age. For female fitness, experimental females were given the opportunity to mate with standard LHm (Larry Harshman, moderate density population) (Rice *et al*. 2005) males for 5 hours at a concentration of 30 flies per vial (1:1 sex-ratio). After mating, females were placed in separate vials to lay eggs for a period of 14 hours. Female fecundity (number of eggs laid - *n* eggs · female^-1^), fertility (adult offspring produced from those eggs - *n* adults · female^-1^) and eggs-adults survival ((adults · eggs^-1^) %) were then measured (Camus *et al*. 2017a; Camus *et al*. 2020b). For male non-competitive fitness, experimental males were given the opportunity to mate with standard flies from the massbred population LHm females for 5 hours at a concentration of 30 flies per vial (1:1 sex-ratio). Females were then sorted in individual vials and allowed to lay eggs for 48 hours. They were then transferred to new vials and left to oviposit for additional 48 hours. Male fertility (adult offspring from the 96 hours lay - *n* adults · female^-1^) was then measured (Camus *et al*. 2020a). For male competitive fitness, a trio of experimental males competed with a trio of LHm *bw-* males (outbred population with homozygous recessive brown eye mutation) for the mating of six LHm *bw-* virgin females, over a period of 24 hours. Red-eyed (wild type - WT) progeny was assigned to the experimental line while brown-eyed progeny to the competitor line. Male fertility was defined as the percentage of red-eyed adults of the total offspring yield ((WT · adults^-1^) %) (Camus *et al*. 2017a).

### Larval development

Experimental flies were placed in separate oviposition chambers and let mate - lay eggs for 2 hours. Eggs were then gently collected and placed in separate vials at a concentration of 30 eggs per vial. All vials were screened for newly eclosed adults three times daily (10 am, 13 pm, 16 pm), for a period of 14 days. This gave ample time for all developing flies to eclose, with any remaining pupae deemed dead. Both development time (hours) and sex were recorded. Survival to adulthood was also measured as the percentage of successfully hatched adults in each vial ((*n* adults · *n* eggs^-1^) %) (Rodríguez *et al*. 2021).

### Thermal tolerance

Heat tolerance assays involved exposing non-virgin flies to a 39°C environment (glass vials immersed in a circulating water bath) and recording the time (min) taken for each fly to succumb to heat stress (heat knock-down) (Hoffmann *et al*. 2002; Camus *et al*. 2017b). Cold tolerance assays involved exposing non-virgin flies to a 0°C environment (plastic tubes immersed in an ice-slurry water bath) for 4 hours to induce chill coma response. Tubes were then placed at 25°C and the time (min) taken for each fly to regain consciousness (standing upright) recorded (CCRT - chill-coma recovery time). Cold tolerance was expressed as 120 minus CCRT (Camus *et al*. 2017b).

### Data analysis

Data were analysed with the software R (R Core Team 2021) and several supporting packages (see supporting information). Respirometry data was further analysed using principal components analysis. Metabolic continuous variables were centered and standardized prior analysis, principal components were then extracted and analyzed as single parameters. A linear mixed model was implemented for each parameter, considering mitochondrial haplotype (*‘mtDNA’*), nuclear background (*‘nDNA’*) and sex (*‘sex’*) as categorical fixed effects, as well as their two- and three-way interactions. Models accounted for differences in population replicate(*‘pop’*), generational sampling block (*‘batch’*), fly age (*‘day’*), trial (*‘run’*) and vial (*‘vial’*), which were included as random effects. A generalized linear mixed model fitting the same fixed and random effects was implemented for traits following either Poisson or binomial distribution. The significance of the three main factors and their possible interactions were determined through a Type III ANOVA, followed by *post hoc* multi comparison and Hommel adjustment for multiple testing. The best fitting model was determined through a step-wise simplification by backward elimination of nonsignificant highest order effects. Detailed information is provided in supplementary material and tables s1-s11.

## Results

### Mitochondrial phenotype

The impact of mitonuclear interactions was tested at the level of mitochondrial respiration (O2 fluxes normalized for tissue mass - pmol O2 · s^−1^ · mg^−1^) sustained by different combination of substrates. An interaction effect between the mitochondrial and the nuclear genomes was revealed for the max coupled (state 3) respiration (CI+ProDH+CII+GpDHP, *F*=6.59, *P*=0.014*), max uncoupled (state 3u) respiration (CI+ProDH+CII+GpDHE, *F*=4.66, *P*=0.037*), and cytochrome *c* oxidase standalone capacity (CIVE, *F*=9.12, *P*=0.0044**) (figures 1A,B, s2A, table s1). Overall, these results indicate that mitonuclear interactions can have an impact on mitochondrial respiration across both sexes, with mismatched cybrids (‘mT’, ‘tM’) showing significant increased respiratory rates compared to the native matched line ‘tT’, as well as a trend of increased respiration with respect to the other native matched line ‘mM’ (table s1). These results were also supported by the analysis of the principal components (figure s3, table s1), where an interaction effect between mtDNA and nDNA reflecting the cybrid-specific mitochondrial phenotype was found for PC1 (*F*=5.74, *P*=0.02*), with CI+ProDH+CII+GpDHP, CI+ProDH+CII+GpDHE and CIVE parameters highly loading on this axis (figure s3C,E). Variations in respiratory rates among mitonuclear lines did not associate with changes in tissue mass and mtDNA content. Thorax weight was influenced by sex (*F*=77.72, *P*=3.47e-11***), while mtDNA content by a nDNA by sex interaction (*F*=6.73, *P*=0.011*) (figures s2B,C, tables s1, s2), suggesting that males have lower body mass and higher mtDNA content than females, the latter only in lines with Townsville nuclear background (‘tT’, ‘mT’).

**Figure 1.**
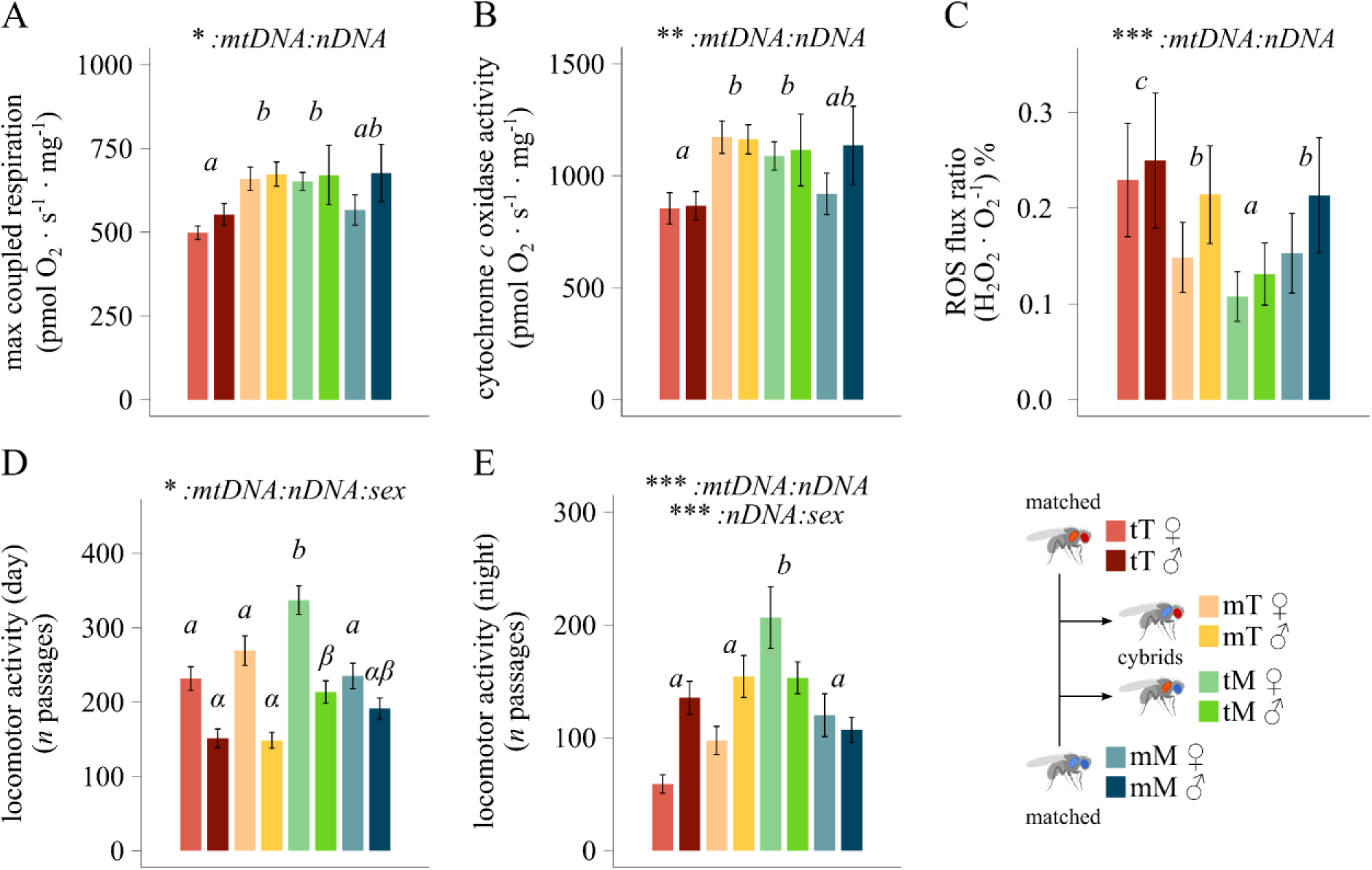
Mitochondrial phenotype and locomotor activity across mitonuclear lines and sexes. **(A-B)** Mitochondrial respiration in permeabilized fly thoraces (pmol O2 · s^-1^ · mg^-1^) reflecting the *(a)* max coupled respiration sustained by CI, CII, ProDH and GpDH-linked substrates, and *(b)* cytochrome *c* oxidase standalone capacity (*n*=6). **(C)** Hydrogen peroxide production over oxygen consumption during max coupled respiration ((H2O2 · O2^-1^) %) (*n*=6). **(D-E)** Fly locomotor activity during *(d)* daytime and *(e)* night (*n* passages) (*n*=62-64). Statistical analyses: linear mixed model; Fixed effects: *’mtDNA’*, *‘nDNA’* and *‘sex’*, plus their interactions. Significance was determined by means of a type III ANOVA. Letters indicate statistical difference following a *post-hoc* multi comparison test. Comparison in (D) was run separately for each sex. Data shown as mean ± sem. \**p* ≤ 0.05; \*\**p* ≤ 0.01; \*\*\**p* ≤ 0.001. A detailed summary is reported in supplementary tables s1, s3, s4.

The analysis of flux control ratios (FCR) (i.e. qualitative analysis with parameters normalized for their own max uncoupled respiration), as well as flux control factors (FCF), revealed no overall differences in substrate preferences dictated by mitonuclear combination (figure s4, table s1). That said, FCR CIVE (figure s4A) and CIV excess capacity (jExCIV) (figure s4B) were influenced by mitonuclear combination, with ‘mT’ cybrid showing a trend of higher activity compared to all other lines. A main effect of sex was revealed for CIP (*F*=4.51, *P*=0.039*), CI+ProDHP (*F*=5.97, *P*=0.018*) and CI+ProDH+CIIP (*F*=8.81, *P*=0.0049**) expressed as FCR (figure s4A), for CIL (*F*=10.57, *P*=0.0022**), CI+ProDHP (*F*=4.57, *P*=0.039*) and CI+ProDH+CIIP (*F*=6.26, *P*=0.017*) expressed as O2 fluxes (figure s2A), as well as for PC2, with CIP, CI+ProDHP and CI+ProDH+CIIP highly loading on it (*F*=4.52, *P*=0.039*) (figure s3D,F). Differences dictated by sex were also revealed for the FCF G3PCF, reflecting the increase in respiration following glycerophosphate addition (figure s4B). Overall, these results suggest sex-specific differences in substrate-preference, with males having higher respiratory rates sustained by CI, ProDH and CII complexes, while females relying more on GpDH activity to sustain maximal state 3 respiration.

Reactive oxygen species production rate (pmol H2O2 · s^−1^ · mg^−1^) was measured in parallel with mitochondrial respiration (table s3). Differences among sexes were revealed in ROS fluxes during max coupled respiration and during total inhibition of the ETS (maximal ROS production) (*F*=10.16, *P*=0.0028** and *F*=31, *P*=1.754e-06***, respectively), with males having generally higher ROS production rates compared to females (figures s5A,B). When further scrutinizing ROS efflux rate over the concomitant respiratory rates (H2O2 · O2^-1^), we however found a significant mtDNA by nDNA interaction during max coupled respiration (*F*=13.18, *P*=0.0009***), indicating lower ROS production per molecule of oxygen consumed in mismatched cybrids (‘mT’, ‘tM’) compared to their genetically closest (at the level of nuclear background) matched line (‘tT’, ‘mM’) (figure 1C, table s3). The max ROS ratio (inhibited ETS) was mainly influenced by both nDNA and sex, without interaction (figure s5C).

### Life-history traits

In addition to mitochondrial physiology, the impact of mitonuclear interactions was further tested on different life-history traits. An interaction effect between the mitochondrial and the nuclear genome was found for fly locomotor activity during the day, which also varied across sexes (*F*=5.02, *P*=0.025*), and during the night (*F*=21.27, *P*=5.056e-06***) (figures 1D,E, s6, table s4). In line with respirometry results, ‘tM’ cybrids are more active than both ‘tT’ and ‘mM’ parental lines during both day and night, whereas the activity of ‘mT’ cybrids does not significantly differ. That said, a higher night locomotor activity of cybrids compared to the closest parental line is supported when comparing lines within a common nuclear background (‘tT’-‘mT’ and ‘tM’-‘mM’) (table s4).

Mitonuclear interactions were also found to have a pervasive effect on reproductive success and offspring development. Mitonuclear interactions impacted female fecundity (*X^2^*=58, *P*=2.611e-14***), fertility (*X^2^*=99.79, *P*<2.2e-16***), and survival (*X^2^*=65.60, *P*=5.510e-16***), with cybrid lines laying fewer eggs and having less offspring compared to both parental populations (figures 2A,B, table s5). Mitonuclear hybrids additionally displayed reduced egg to adult survival compared to ‘tT’ matched line, whereas only a trend was found when comparing these to the ‘mM’ line (figure s7A, table s5). Differences dictated by the mitonuclear genotype existed also for male fertility in a non-competitive environment (*X^2^*=14.31, *P*=0.00015***), with cybrids having fewer offspring than ‘tT’ parental population, although not for ‘mM’ (figure 2C, table s6). On the other hand, male fertility in a competitive environment was influenced by both the mtDNA and the nDNA, without interaction (figure s7B, table s7). Larval developmental time was also influenced by the mitonuclear combination (*F*=58.21, *P*=3.890e-14***), with female and male individuals of the mismatched ‘tM’ mitonuclear line developing slower than flies from both parental populations (figure 2D, table s8). Although individuals from ‘mT’ mismatched line only display a trend of slower developmental rate compared to their closest parental population (‘tT’), a significant difference between the two lines was revealed in females when testing the impact of mtDNA within nuclear background and sex (table s8). The interaction between mtDNA and nDNA genomes additionally influenced the survival rate during development (*X^2^*=17.06, *P*=3.621e-05***), with individuals from Townsville parental line (‘tT’) better surviving to adulthood compared to both cybrids (‘mT’ and ‘tM’) and Melbourne parental line (‘mM’) (figure 2E, table s9).

**Figure 2.**
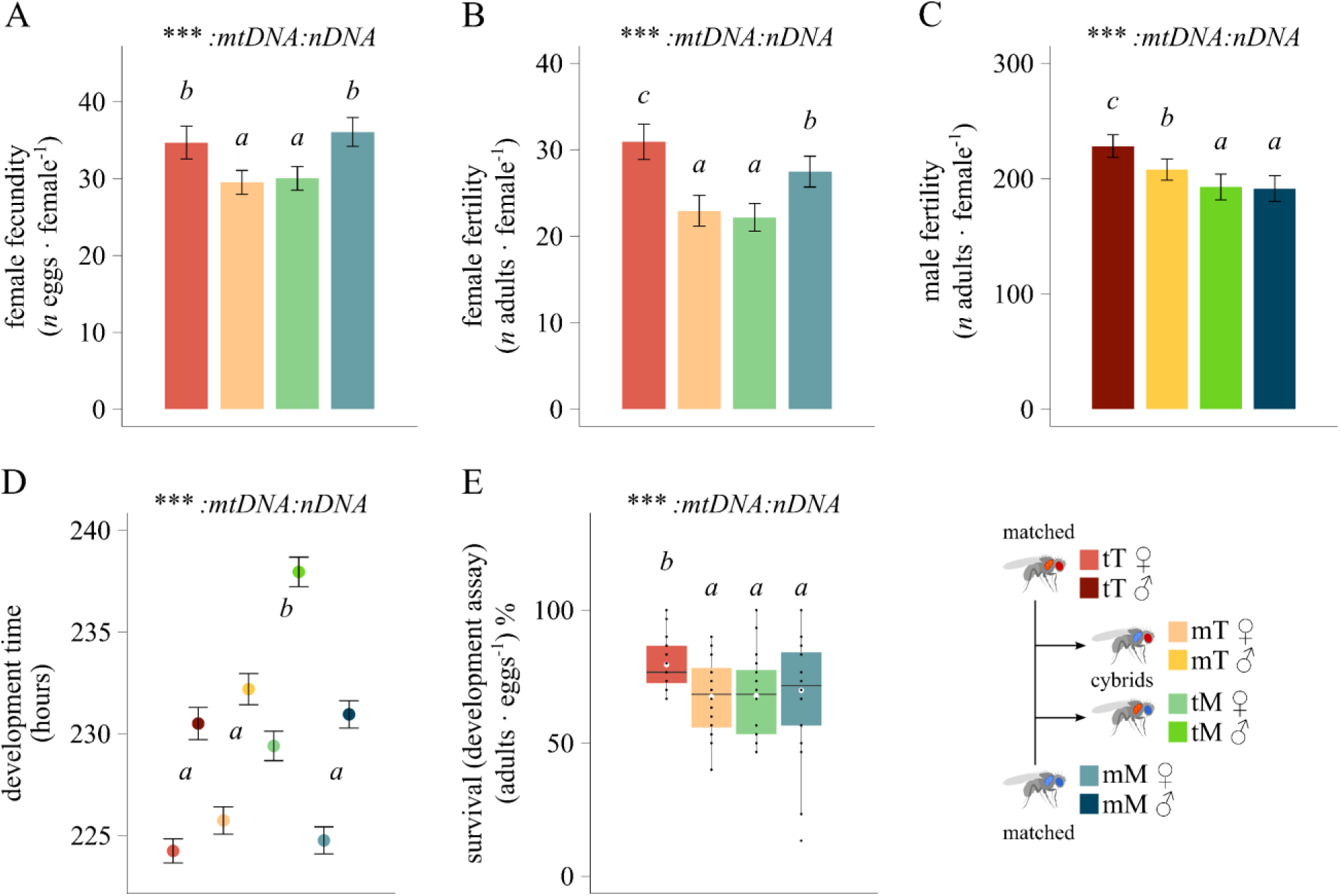
Reproductive fitness and development time across mitonuclear genotypes and sexes. **(A-B)** Female reproductive success expressed by *(a)* the number of eggs laid by focal females mated with standardized LHm males (fecundity - *n* eggs · female^-1^), and *(b)* the number of adult offspring produced for each lay (fertility - *n* adults · female^-1^) (*n*=56-60). **(C)** Male reproductive success expressed as the number of offspring produced when standard LHm females mated with focal males of each mitonuclear line (fertility - *n* adults · female^-1^) (*n*=37-39). **(D-E)** Developmental fitness expressed by the egg-adult *(d)* development time (hours) (*n*=177-280), *(e)* percentage of adults survival in each vial ((adults · eggs^-1^) %) (*n*=20). Statistical analyses: (A, B, C, E) generalized linear mixed model; (D) linear mixed model. Fixed effects: *’mtDNA’*, *‘nDNA’* and *‘sex’* (only for D), plus their interactions. Significance was determined by means of a type III ANOVA. Letters indicate statistical difference following a *post-hoc* multi comparison test. Data (A-D) shown as mean ± sem. \**p* ≤ 0.05; \*\**p* ≤ 0.01; \*\*\**p* ≤ 0.001. A detailed summary is reported in supplementary tables s5, s6, s8, s9.

### Thermal tolerance

We finally assessed the influence of nuclear and mitonuclear genomes on fly thermal performance. No interaction effects between mtDNA, nDNA and sex were detected for either heat and cold tolerance parameters, which were influenced by the solely nDNA (*F*=30.48, *P*=6.102e-08*** and *F*=19.87, *P*=1.033e-05***, respectively) and sex (*F*=56.56, *P*=3.695e-13*** and *F*=48.57, *P*=1.073e-11***, respectively) (tables s10, s11). In line with the natural latitudinal segregation of the two native populations, mitonuclear combinations with northern-derived Townsville nDNA (‘tT’, ‘mT’) resisted more to heat shock than flies bearing the southern-derived Melbourne nDNA (‘tM’, ‘mM’) (figure s8A). On the other hand, lines with Melbourne nDNA better recovered from cold shock than lines with Townsville nDNA (figure s8B). Females and males also showed divergent thermal tolerance patterns, with female flies better able to withstand cold stress than males, whereas male flies being more resistant to heat stress than their female counterparts (figures s8A,B, tables s10, s11).

## Discussion

One of the many consequences of local adaptation is the divergence of genomic information. This information not only applies to the nuclear genome, as recent work has highlighted the role of the mitochondrial genome as an important source of adaptive variation (Blier *et al*. 2001; Dowling *et al*. 2008; Lane 2009, 2011; Wolff *et al*. 2014). Here, we explored how mitonuclear epistasis shapes energy flow at the cellular level, and how variation in aerobic metabolism might influence key evolutionary and ecological patterns, from life-history phenotypes to adaptation. We investigated this question using a *Drosophila melanogaster* mitonuclear panel composed of coadapted populations and mitonuclear decoupled cybrid populations. Our results support the notion that introgression of naturally-occurring mtDNA can have profound repercussions at different levels of biological hierarchy, from mitochondrial function to organismal fitness, with potential impacts on population ecology.

We observed higher overall respiratory rates in *Drosophila* cybrids compared with coevolved populations, in both females and males (figures 1A,B, s2A, s3), linked with a decreased amount of ROS production per molecule of oxygen consumed (figure 1C). Curiously, these differences were not underpinned by changes in thorax mass or mitochondrial content (figures s2B,C), nor with changes in substrate preferences, which could indicate a shift in metabolic flux to compensate for the genomic mismatch (figure s4). The genetic basis of the phenotypic differences observed is underpinned by 15 synonymous mutations SNP, which are widespread among most coding genes, including *ND2*, *ND6*, *CYTB*, *COX1*, *COX2*, *COX3*, *ATP6*, mt-*tRNAE* and mt-*sRNA* (1 mutation each), as well as *ND1*, *ND4* and *ND5* (2 mutations each) (Camus *et al*. 2017b). Although traditionally considered functionally silent, synonymous mutations can indeed be evolutionary significant, impacting mRNA stability, translational speed, folding and post-translational modifications of proteins, as well as enzyme structure and function (Hurst 2011; Shabalina *et al*. 2013; Jiang *et al*. 2022). This is also the case of variations in both control region and alternative genes of the mitochondrial genome which could impact the phenotype (Lee *et al*. 2015; Rollins *et al*. 2016; Hopkins *et al*. 2017; Breton 2021; Pozzi & Dowling 2021; Kienzle *et al*. 2023). In the case of our two main haplotypes (‘t’ and ‘m’), previous work highlighted differences in gene expression patterns between them against a common foreign nuclear background (Camus *et al*. 2017b). It is therefore possible (though as yet unproved in our massbred populations), that a mismatch-induced change in the transcriptional rate, and by extension the number of ETS enzymes, might underpin the cybrid-specific mitochondrial phenotype described here.

Hybridisation can sometimes be beneficial, providing an effective source of adaptive alleles that overcome the fitness cost of mitonuclear incompatibility (Hill 2019). This has been exemplified in both migrating birds and *Drosophila* showing signs of adaptive mitonuclear co-introgression (Toews et al. 2014; Beck et al. 2015; Morales et al. 2018). That said, the general pattern expected from mitonuclear disfunction is a breakdown of mitochondrial efficiency (reduced OXPHOS capacity), associated with deleterious fitness consequences (Burton *et al*. 2006; Ellison *et al*. 2008; Meiklejohn *et al*. 2013). The question therefore arises on whether the increase in cybrid respiratory rate described here could underpin improved OXPHOS efficiency (ATP/O ratio), or rather reflect an overall dysregulation of mitochondrial function via some form of overcompensation for deficient mitochondria (Moreno-Loshuertos *et al*. 2006). We did not look specifically at ATP production, but our results for leak respiration (CIL), coupling efficiency (jP) and ROS flux (figures 1C, s4, s5) all suggest that the proportion of energy dispersed as ROS production or futile proton cycling (leak respiration) does not increase because of the high mitochondrial respiration of cybrids. This provides indirect support for normally functional OXPHOS in cybrid populations. Further support comes from the analysis of locomotor activity, a generally high-energy demanding process (likely requiring healthy mitochondria), which was also maintained, even increased in ‘tM’ cybrids (figures 1D,E, s6). However, despite the increase in cybrid respiration, these were generally less fertile than coevolved populations, implying that there is indeed a life-history cost to mismatching mitonuclear genomes. Notably, cybrids laid fewer eggs (figure 2A), had a slower larval development (figure 2D), reduced egg-adult survival (figures 2E, s7A) and decreased number of offspring (figures 2B,C).

Organisms must adopt life-history strategies when allocating the finite energetic resources available, as increasing the energy allocation to one trait inevitably reduces the availability for the others (Chang et al., 2021). Examples of trade-offs between locomotory metabolic processes, (locomotor activity or flight), and mainly biosynthetic ones (reproduction and growth) are widespread in literature (Zhang *et al*. 2009; Gibbs *et al*. 2010; Husak *et al*. 2016; Chang *et al*. 2021). Although high metabolism generally correlates with fast development, early reproduction and decreased longevity (Pettersen *et al*. 2016), previous evidence in wild-type *Drosophila* lines linked high complex IV activity and metabolic rates with lowered fecundity and lifespan (Ballard *et al*. 2007a; Mołoń *et al*. 2020). Furthermore, mitonuclear interactions were found to have a substantial impact on resource allocation and life-history trade-offs in flies (Camus *et al*. 2020b). Our results support mitonuclear effects on *Drosophila* phenotype, involving some sort of trade-off between mitochondrial bioenergetics and organismal fitness in cybrid populations.

The physiological basis of these life-history dynamics are uncertain, as our measures of mitochondrial bioenergetics were obtained from thoraces (flight muscle), whose bioenergetic requirements are likely to be very different to tissues more heavily involved with reproduction. Compared to somatic tissues, gonads primarily need to power biosynthesis for gamete production and the relative requirements for ATP synthesis are several fold lower than flight muscle (Wetzker & Reinhardt 2019; Camus *et al*. 2023). The different metabolic profiles of somatic and reproductive tissues are also accompanied, at least in some mammals, by the expression of gamete-specific nuclear isoforms of some OXPHOS genes (Huttemann *et al*. 2003; Liu *et al*. 2006), opening up the opportunity for divergent mitonuclear coevolution (and incompatibilities) in somatic tissues and sex organs. Given these different energetic requirements between tissues, the trade-off in our cybrid flies might be explained by the way that mitochondria power biosynthesis rather than OXPHOS itself. At the level of gonads, this may result in the compromised fertility observed in cybrids, as well as underpinning the slower growth and decreased survival of their larvae. Reallocation of resources away from gamete production (and growth in general) could in turn explain the higher aerobic capacity of somatic tissues and locomotor activity of cybrids. Our future studies will aim to investigate whether the main phenotypic effect of mitonuclear mismatch might differ in gonads, potentially explaining the decrease in components of fitness.

The colonisation of Australia by *D. melanogaster* is quite recent and traces back just a few hundreds of years (Hoffmann & Weeks 2007; Adrion *et al*. 2015). Despite continued admixture and gene-flow between populations (Bergland *et al*. 2016), the Australian eastern cline has remained stable over the decades; defined by a predictable variation in phenotype and genotype suggesting ongoing climatic selection (Hoffmann & Weeks 2007; Sgro *et al*. 2010; Adrion *et al*. 2015; Camus *et al*. 2017b; Lajbner *et al*. 2018; Chakraborty *et al*. 2020). Similar to previous evidence, we detected differences in thermal tolerance between our two genomic backgrounds, with Townsville populations and Melbourne populations respectively being more resistant to heat and cold stress (figure s8). Temperature is a well-known metabolic stressors, which could exacerbate, and even drive, the main effect of mitochondrial introgression (Rank *et al*. 2020). Our data highlights the nuclear genome as main contributor to thermal tolerance, with the mtDNA having a much lower contribution than previously measured (Camus *et al*. 2017b; Lajbner *et al*. 2018). However, in these studies, the mtDNA variants were coupled to a standardised isogenic nuclear background (Camus *et al*. 2017b), allowing for a more precise control of genetic effects. It is possible that mtDNA effects on thermal traits are more subtle and get swamped by the large genetic variance coming from the nuclear genome. Furthermore, we cannot exclude that adaptation to chronic increased temperature (or vast temperature fluctuations) might have a mitonuclear contribution, exacerbating the main phenotypic effect of mitonuclear incompatibility herein described. Future research will therefore aim to test that.

Mitonuclear incongruences have long been proposed as specific cases of Bateson– Dobzhansky–Muller incompatibilities (Burton *et al*. 2006). Depending on the severity of the incompatibility (either via high genetic divergence or large effect SNPs), mitonuclear epistasis may restrict gene flow between populations and has the potential to reinforce reproductive isolation (Gershoni *et al*. 2009; Burton & Barreto 2012; Hill 2017, 2019; Burton 2022). Regardless the absence of allopatry and the low mitochondrial genetic divergence between fly populations, we found evidence that naturally-occurring mitonuclear disruptions strongly decreased reproductive performance in both female and male cybrids. These results support the existence of partial genetic barriers dictated by the mitonuclear combination. Consequently, mitonuclear coadaptation under climatic selection might contribute to the evolutionary trajectory, clinal distribution and future ecological adaptation of natural populations.

In this study we tested the extent to which mitonuclear epistasis might impact organismal fitness and population persistence in divergent, locally-adapted insect populations. Our results provide clear evidence that even small differences in mitonuclear genotype can impair both fecundity and male fertility, potentially reducing gene flow among fly populations locally-adapted to different thermal niches along a latitudinal cline. Curiously, these mitonuclear genotypes apparently boosted respiratory outputs and locomotor activity, suggesting a possible trade-off between aerobic capacity and fertility, which might reflect limited metabolic plasticity in mitochondrial function. We live in a rapidly changing world, in which natural populations are experiencing unprecedented changes in temperature, diet and geographical distribution. If mitonuclear interactions constraint metabolic plasticity and fitness, then it will be important to include mitonuclear epistasis in ecological studies of the adaptive capacity of natural populations.

## Supporting information

Supplementary tables and stats

## AKNOWLEDGEMENTS

This project has received funding from the European Union’s Horizon 2020 research and innovation programme under the Marie Skłodowska-Curie grant agreement No. 101030803 to S.B. M.F.C was funded by a UKRI Fellowship (NE/V014307/1).

## Notes

https://doi.org/10.6084/m9.figshare.24162879

### Competing Interest Statement

The authors have declared no competing interest.

## REFERENCES

1. Adrion, J.R., Hahn, M.W. & Cooper, B.S. (2015). Revisiting classic clines in Drosophila melanogaster in the age of genomics. Trends Genet, 31, 434–444.

2. Anderson, L., Camus, M.F., Monteith, K.M., Salminen, T.S. & Vale, P.F. (2022). Variation in mitochondrial DNA affects locomotor activity and sleep in Drosophila melanogaster. Heredity, 129, 225–232.

3. Ballard, J.W., Melvin, R.G., Katewa, S.D. & Maas, K. (2007a). Mitochondrial DNA variation is associated with measurable differences in life-history traits and mitochondrial metabolism in Drosophila simulans. Evolution, 61, 1735–1747.

4. Ballard, J.W., Melvin, R.G., Miller, J.T. & Katewa, S.D. (2007b). Sex differences in survival and mitochondrial bioenergetics during aging in Drosophila. Aging cell, 6, 699–708.

5. Barreto, F.S., Watson, E.T., Lima, T.G., Willett, C.S., Edmands, S., Li, W. et al. (2018). Genomic signatures of mitonuclear coevolution across populations of *Tigriopus californicus*. Nature Ecology & Evolution, 2, 1250–1257.

6. Barrientos, A., Kenyon, L. & Moraes, C.T. (1998). Human Xenomitochondrial Cybrids: cellular models of mitochondrial complex I deficiency. Journal of Biological Chemistry, 273, 14210–14217.

7. Bergland, A.O., Tobler, R., González, J., Schmidt, P. & Petrov, D. (2016). Secondary contact and local adaptation contribute to genome-wide patterns of clinal variation in Drosophila melanogaster. Molecular Ecology, 25, 1157–1174.

8. Bettinazzi, S., Rodríguez, E., Milani, L., Blier, P.U. & Breton, S. (2019). Metabolic remodelling associated with mtDNA: insights into the adaptive value of doubly uniparental inheritance of mitochondria. Proceedings of the Royal Society B: Biological Sciences, 286, 20182708.

9. Biot-Pelletier, D., Bettinazzi, S., Gagnon-Arsenault, I., Dubé, A.K., Bédard, C., Nguyen, T.H.M. et al. (2023). Evolutionary Trajectories are Contingent on Mitonuclear Interactions. Molecular Biology and Evolution, 40.

10. Blier, P.U., Dufresne, F. & Burton, R.S. (2001). Natural selection and the evolution of mtDNA-encoded peptides: evidence for intergenomic co-adaptation. Trends Genet, 17, 400–406.

11. Breton, S. (2021). Mitochondrial Russian doll genes may explain some discrepancies in links between mtDNA mutations and mitochondrial diseases. BioEssays, 43, 2100104.

12. Burger, J.R., Hou, C. & Brown, J.H. (2019). Toward a metabolic theory of life history. Proceedings of the National Academy of Sciences, 116, 26653–26661.

13. Burton, R.S. (2022). The role of mitonuclear incompatibilities in allopatric speciation. Cellular and Molecular Life Sciences, 79, 103.

14. Burton, R.S. & Barreto, F.S. (2012). A disproportionate role for mtDNA in Dobzhansky-Muller incompatibilities? Mol Ecol, 21, 4942–4957.

15. Burton, R.S., Ellison, C.K. & Harrison, J.S. (2006). The sorry state of F2 hybrids: consequences of rapid mitochondrial DNA evolution in allopatric populations. The American naturalist, 168 Suppl 6, S14–24.

16. Camus, M.F., Fowler, K., Piper Matthew, W.D. & Reuter, M. (2017a). Sex and genotype effects on nutrient-dependent fitness landscapes in *Drosophila melanogaster*. Proceedings of the Royal Society B: Biological Sciences, 284, 20172237.

17. Camus, M.F., Moore, J. & Reuter, M. (2020a). Nutritional geometry of mitochondrial genetic effects on male fertility. Biology Letters, 16, 20190891.

18. Camus, M.F., O’Leary, M., Reuter, M. & Lane, N. (2020b). Impact of mitonuclear interactions on life-history responses to diet. Philos Trans R Soc Lond B Biol Sci, 375, 20190416.

19. Camus, M.F., Rodriguez, E., Kotiadis, V., Carter, H. & Lane, N. (2023). Redox stress shortens lifespan through suppression of respiratory complex I in flies with mitonuclear incompatibilities. Experimental Gerontology, 175, 112158.

20. Camus, M.F., Wolff, J.N., Sgro, C.M. & Dowling, D.K. (2017b). Experimental Support That Natural Selection Has Shaped the Latitudinal Distribution of Mitochondrial Haplotypes in Australian *Drosophila melanogaster*. Mol Biol Evol, 34, 2600–2612.

21. Chakraborty, A., Sgrò, C.M. & Mirth, C.K. (2020). Does local adaptation along a latitudinal cline shape plastic responses to combined thermal and nutritional stress? Evolution, 74, 2073–2087.

22. Chang, H., Guo, X., Guo, S., Yang, N. & Huang, Y. (2021). Trade-off between flight capability and reproduction in Acridoidea (Insecta: Orthoptera). Ecol Evol, 11, 16849–16861.

23. Chapdelaine, V., Bettinazzi, S., Breton, S. & Angers, B. (2020). Effects of mitonuclear combination and thermal acclimation on the energetic phenotype. Journal of Experimental Zoology Part A: Ecological and Integrative Physiology, 333, 264–270.

24. Deremiens, L., Schwartz, L., Angers, A., Glémet, H. & Angers, B. (2015). Interactions between nuclear genes and a foreign mitochondrial genome in the redbelly dace Chrosomus eos. Comparative Biochemistry and Physiology Part B: Biochemistry and Molecular Biology, 189, 80–86.

25. Dey, R., Barrientos, A. & Moraes, C.T. (2000). Functional Constraints of Nuclear-Mitochondrial DNA Interactions in Xenomitochondrial Rodent Cell Lines. Journal of Biological Chemistry, 275, 31520–31527.

26. Dowling, D.K., Friberg, U. & Lindell, J. (2008). Evolutionary implications of non-neutral mitochondrial genetic variation. Trends Ecol Evol, 23, 546–554.

27. Ellison, C.K. & Burton, R.S. (2006). Disruption of mitochondrial function in interpopulation hybrids of *Tigriopus californicus*. Evolution, 60, 1382–1391.

28. Ellison, C.K., Niehuis, O. & Gadau, J. (2008). Hybrid breakdown and mitochondrial dysfunction in hybrids of Nasonia parasitoid wasps. J Evol Biol, 21, 1844–1851.

29. Gershoni, M., Templeton, A.R. & Mishmar, D. (2009). Mitochondrial bioenergetics as a major motive force of speciation. Bioessays, 31, 642–650.

30. Gibbs, M., Breuker, C.J., Hesketh, H., Hails, R.S. & Van Dyck, H. (2010). Maternal effects, flight versus fecundity trade-offs, and offspring immune defence in the Speckled Wood butterfly, Pararge aegeria. BMC Evolutionary Biology, 10, 345.

31. Gnaiger, E. (2020). Mitochondrial Pathways and Respiratory Control An Introduction to OXPHOS Analysis, 5th Edition. Bioenergetics Communications.

32. Healy, T.M. & Burton, R.S. (2020). Strong selective effects of mitochondrial DNA on the nuclear genome. Proc Natl Acad Sci U S A, 117, 6616–6621.

33. Hill, G.E. (2017). The mitonuclear compatibility species concept. The Auk, 134, 393–409.

34. Hill, G.E. (2019). Reconciling the Mitonuclear Compatibility Species Concept with Rampant Mitochondrial Introgression. Integrative and Comparative Biology, 59, 912–924.

35. Hill, G.E. (2020). Mitonuclear Compensatory Coevolution. Trends Genet, 36, 403–414.

36. Hoffmann, A.A., Anderson, A. & Hallas, R. (2002). Opposing clines for high and low temperature resistance in Drosophila melanogaster. Ecology Letters, 5, 614–618.

37. Hoffmann, A.A. & Weeks, A.R. (2007). Climatic selection on genes and traits after a 100 year-old invasion: a critical look at the temperate-tropical clines in Drosophila melanogaster from eastern Australia. Genetica, 129, 133–147.

38. Hopkins, J.F., Sabelnykova, V.Y., Weischenfeldt, J., Simon, R., Aguiar, J.A., Alkallas, R. et al. (2017). Mitochondrial mutations drive prostate cancer aggression. Nature Communications, 8, 656.

39. Hurst, L.D. (2011). The sound of silence. Nature, 471, 582–583.

40. Husak, J.F., Ferguson, H.A. & Lovern, M.B. (2016). Trade-offs among locomotor performance, reproduction and immunity in lizards. Functional Ecology, 30, 1665–1674.

41. Huttemann, M., Jaradat, S. & Grossman, L.I. (2003). Cytochrome *c* oxidase of mammals contains a testes-specific isoform of subunit VIb--the counterpart to testes-specific cytochrome *c*? Molecular reproduction and development, 66, 8–16.

42. Jiang, Y., Neti, S.S., Sitarik, I., Pradhan, P., To, P., Xia, Y. et al. (2022). How synonymous mutations alter enzyme structure and function over long timescales. Nature Chemistry.

43. Kienzle, L., Bettinazzi, S., Choquette, T., Brunet, M., Khorami, H.H., Jacques, J.-F. et al. (2023). A small protein coded within the mitochondrial canonical gene nd4 regulates mitochondrial bioenergetics. BMC Biology, 21, 111.

44. Lajbner, Z., Pnini, R., Camus, M.F., Miller, J. & Dowling, D.K. (2018). Experimental evidence that thermal selection shapes mitochondrial genome evolution. Sci Rep, 8, 9500.

45. Lane, N. (2009). Biodiversity: On the origin of bar codes. Nature, 462, 272–274.

46. Lane, N. (2011). Mitonuclear match: optimizing fitness and fertility over generations drives ageing within generations. Bioessays, 33, 860–869.

47. Lee, C., Zeng, J., Drew, Brian G., Sallam, T., Martin-Montalvo, A., Wan, J. et al. (2015). The Mitochondrial-Derived Peptide MOTS-c Promotes Metabolic Homeostasis and Reduces Obesity and Insulin Resistance. Cell Metabolism, 21, 443–454.

48. Lee, H.Y., Chou, J.Y., Cheong, L., Chang, N.H., Yang, S.Y. & Leu, J.Y. (2008). Incompatibility of nuclear and mitochondrial genomes causes hybrid sterility between two yeast species. Cell, 135, 1065–1073.

49. Liu, Z., Lin, H., Ye, S., Liu, Q.-y., Meng, Z., Zhang, C.-m., et al. (2006). Remarkably high activities of testicular cytochrome *c* in destroying reactive oxygen species and in triggering apoptosis. Proceedings of the National Academy of Sciences of the United States of America, 103, 8965–8970.

50. Ma, H., Marti Gutierrez, N., Morey, R., Van Dyken, C., Kang, E., Hayama, T., et al. (2016). Incompatibility between Nuclear and Mitochondrial Genomes Contributes to an Interspecies Reproductive Barrier. Cell Metabolism, 24, 283–294.

51. Meiklejohn, C.D., Holmbeck, M.A., Siddiq, M.A., Abt, D.N., Rand, D.M. & Montooth, K.L. (2013). An Incompatibility between a Mitochondrial tRNA and Its Nuclear-Encoded tRNA Synthetase Compromises Development and Fitness in Drosophila. PLOS Genetics, 9, e1003238.

52. Mołoń, M., Dampc, J., Kula-Maximenko, M., Zebrowski, J., Mołoń, A., Dobler, R. et al. (2020). Effects of Temperature on Lifespan of Drosophila melanogaster from Different Genetic Backgrounds: Links between Metabolic Rate and Longevity. Insects, 11.

53. Morales, H.E., Pavlova, A., Amos, N., Major, R., Kilian, A., Greening, C. et al. (2018). Concordant divergence of mitogenomes and a mitonuclear gene cluster in bird lineages inhabiting different climates. Nature Ecology & Evolution, 2, 1258–1267.

54. Moreno-Loshuertos, R., Acin-Perez, R., Fernandez-Silva, P., Movilla, N., Perez-Martos, A., Rodriguez de Cordoba, S., et al. (2006). Differences in reactive oxygen species production explain the phenotypes associated with common mouse mitochondrial DNA variants. Nat Genet, 38, 1261–1268.

55. Outhwaite, C.L., McCann, P. & Newbold, T. (2022). Agriculture and climate change are reshaping insect biodiversity worldwide. Nature, 605, 97–102.

56. Pettersen, A.K., White, C.R. & Marshall, D.J. (2016). Metabolic rate covaries with fitness and the pace of the life history in the field. Proc Biol Sci, 283.

57. Pozzi, A. & Dowling, D.K. (2021). Small mitochondrial RNAs as mediators of nuclear gene regulation, and potential implications for human health. Bioessays, 43, e2000265.

58. R Core Team (2021). R: A language and environment for statistical computing. R Foundation for Statistical Computing, Vienna, Austria.

59. Rank, N.E., Mardulyn, P., Heidl, S.J., Roberts, K.T., Zavala, N.A., Smiley, J.T. et al. (2020). Mitonuclear mismatch alters performance and reproductive success in naturally introgressed populations of a montane leaf beetle. Evolution, 74, 1724–1740.

60. Rice, W.R., Linder, J.E., Friberg, U., Lew, T.A., Morrow, E.H. & Stewart, A.D. (2005). Inter-locus antagonistic coevolution as an engine of speciation: assessment with hemiclonal analysis. Proc Natl Acad Sci U S A, 102 Suppl 1, 6527–6534.

61. Rodríguez, E., Bettinazzi, S., Inwongwan, S., Camus, M.F. & Lane, N. (2023). Harmonising protocols to measure Drosophila respiratory function in mitochondrial preparations. Bioenergetics Communications, 2023.3.

62. Rodríguez, E., Grover Thomas, F., Camus, M.F. & Lane, N. (2021). Mitonuclear Interactions Produce Diverging Responses to Mild Stress in Drosophila Larvae. Frontiers in genetics, 12.

63. Rollins, L.A., Woolnough, A.P., Fanson, B.G., Cummins, M.L., Crowley, T.M., Wilton, A.N. et al. (2016). Selection on Mitochondrial Variants Occurs between and within Individuals in an Expanding Invasion. Molecular Biology and Evolution, 33, 995–1007.

64. Sackton, T.B., Haney, R.A. & Rand, D.M. (2003). Cytonuclear coadaptation in Drosophila: disruption of cytochrome c oxidase activity in backcross genotypes. Evolution, 57, 2315–2325.

65. Sgro, C.M., Overgaard, J., Kristensen, T.N., Mitchel, K.A., Cockerell, F.E. & Hoffmann, A.A. (2010). A comprehensive assessment of geographic variation in heat tolerance and hardening capacity in populations of *Drosophila melanogaster* from eastern Australia. Journal of Evolutionary Biology, 23, 2484–2493.

66. Shabalina, S.A., Spiridonov, N.A. & Kashina, A. (2013). Sounds of silence: synonymous nucleotides as a key to biological regulation and complexity. Nucleic acids research, 41, 2073–2094.

67. Wang, S., Ore, M.J., Mikkelsen, E.K., Lee-Yaw, J., Toews, D.P.L., Rohwer, S. et al. (2021). Signatures of mitonuclear coevolution in a warbler species complex. Nature Communications, 12, 4279.

68. Wetzker, C. & Reinhardt, K. (2019). Distinct metabolic profiles in Drosophila sperm and somatic tissues revealed by two-photon NAD(P)H and FAD autofluorescence lifetime imaging. Scientific Reports, 9, 19534.

69. Wolff, J.N., Ladoukakis, E.D., Enriquez, J.A. & Dowling, D.K. (2014). Mitonuclear interactions: evolutionary consequences over multiple biological scales. Philos Trans R Soc Lond B Biol Sci, 369, 20130443.

70. Zhang, C., Montooth, K.L. & Calvi, B.R. (2017). Incompatibility between mitochondrial and nuclear genomes during oogenesis results in ovarian failure and embryonic lethality. Development, 144, 2490–2503.

71. Zhang, Y., Wu, K., Wyckhuys, K.A.G. & Heimpel, G.E. (2009). Trade-Offs Between Flight and Fecundity in the Soybean Aphid (Hemiptera: Aphididae). Journal of Economic Entomology, 102, 133–138.

72. Zhu, C.-T., Ingelmo, P. & Rand, D.M. (2014). G×G×E for Lifespan in *Drosophila*: Mitochondrial, Nuclear, and Dietary Interactions that Modify Longevity. PLOS Genetics, 10, e1004354.

