## Supplementary text and figures for "Assessing the role of mitonuclear interactions on mitochondrial function and organismal fitness in natural *Drosophila* populations"

Electronic supplementary material

### SUPPORTING INFORMATION

#### Materials and methods

*Sample collection, mitonuclear lines establishment and rearing conditions.*

Previous work on wild *Drosophila* populations on the east coast of Australia revealed strong genetic signatures in both the nDNA and the mtDNA responsible for local adaptation, with northern and southern populations respectively better adapted to either tropical or temperate environments (Hoffmann *et al.* 2002; Hoffmann & Weeks 2007; Sgro *et al.* 2010; Adrion *et al.* 2015; Bergland *et al.* 2016; Camus *et al.* 2017b; Lajbner *et al.* 2018; Chakraborty *et al.* 2020). For this research, northern and southern nuclear backgrounds were sourced from cline-end massbred *Drosophila* populations in early 2021, transferred to Monash University (Australia) and named accordingly to the specific geographic area of collection: Townsville 'T' (latitude: -19.26, longitude 146.81) and Melbourne 'M' (latitude: -37.77, longitude:

144.99). Isofemale lines were first established from both locations using single wild-caught inseminated females, giving rise to F1 lineage from each isofemale line. Massbred populations were created by collecting virgin F2 flies from each isofemale lineage and mating them *en masse* in a bottle. For the Melbourne population, we used 26 isofemale lines, with each line contributing 12 virgin males and 12 virgin females from each line. Townsville massbred population was generated using 35 isofemale lines, with each line contributing 10 virgin males and 10 virgin females. Each population was maintained over 2 bottles with a density of 400 flies per bottle. Northern ‘t’ and southern ‘m’ mitochondrial haplotypes were sourced from a Coffs Harbour (latitude: -30.29, longitude: 153.11) population collected in 2017, and were already expressed alongside a controlled genetic background. Haplotypes ‘t’ and ‘m' respectively refer to haplotypes ‘A1’ and ‘B1’ in previous studies (Camus *et al.* 2017b). These two haplotypes segregate up the Australian east coast at different population frequencies, with the ‘t’ (A1) group-haplotypes being more prevalent at low latitude, and ‘m’ (B1) group-haplotypes being more prevalent at higher latitude (Camus *et al.* 2017b). All fly lines underwent tetracycline treatment (0.3 $mg{\cdot}mL^{-1}$) against the cytoplasmic endosymbiont *Wolbachia*, whose absence was confirmed through PCR.

Starting from the end populations of the Australian eastern cline, a full-factorial mitonuclear panel was generated over 6 generations. All four mitonuclear lines (both matched and mismatched, to avoid potential bias due to technique itself) were established with the use of balancer chromosomes, following well-established chromosome crossing scheme (figure s1) (Clancy 2008; Zhu *et al.* 2014; Camus *et al.* 2017b). The panel followed a two-letter nomenclature, indicating the mitochondrial genome first and the nuclear background second (lower and uppercase letter, respectively). It included two native strains with matched mitonuclear genome (Townsville – tT, Melbourne – mM) and two cybrid lines (mT and tM), where the mitochondrial and the nuclear genome were reciprocally mismatched. Mass-bred native populations were also preserved for introgression purposes. Specifically, males of mass-bred Townsville or Melbourne lines were regularly used to backcross the experimental lines with a correspondent nuclear background (300 males + 150 virgin females, for each line), in order to maintain genetic variation within the populations. All fly lines were split in two independent populations and maintained in bottles at standard laboratory conditions (25°C, 1:1 protein:carbohydrate P:C diet, 12:12 light:dark day cycle). Standard LHm and LHm *bw-* lines were also reared and used for fertility assays. For the experiments, eggs from the main populations were collected and then reared in density-controlled conditions (20 flies per vial). After eclosion, flies were sorted by sex, let acclimate in new food vials and finally assayed when 4-7 days old.

Throughout the project, the mtDNA haplotypes were regularly verified on a subsample of each population. Total DNA was extracted from 3 pooled adults per sample with the DNeasy Blood & Tissue Kit (Qiagen), and quantified with a NanoDrop 2000 spectrophotometer (Thermo Fisher Scientific Inc). Primers were designed to selectively amplify a specific part of the *cox2* gene, for a total amplicon of ~200 bp. Primer forward: $ALST_F$ (5' ACCTTTACGAAATTCCCATCCTCT 3'); Primer reverse: $ALST_R$ (5' ATGATGCACCGTTAGCATGT 3'). DNA was amplified on a Veriti thermocycler (AB Applied Biosystems), using a Phusion® High-Fidelity DNA Polymerase kit (New England BioLabs® Inc). The reaction (50 µL) was carried  in HF Buffer (1X) with 1.5 mM $MgCl_2$, 200 µM DNTPs, 0.4 µM for each primer, 1U Phusion DNA Polymerase and 100 ng DNA sample. Cycle conditions were as follow: initial denaturation at 98°C for 30 s, followed by 25-35 cycles of 98°C for 5-10 s, 45-72°C for 10-30 s and 72°C for 15-30 s / kb, followed by a final extension at 72°C for 5-10 min. PCR product were then verified on a 1% agarose gel with ethidium bromide dye and HyperLadder™ 100bp (Meridian Bioscience), cleaned with Wizard SV Gel and PCR Clean-Up System (Promega), and sent for Sanger sequencing

(Source Biosciences). Sequences were aligned using the MEGA v11 software (Tamura *et al.* 2021) and the two haplotypes (t, m) discriminated based on known polymorphisms within the *cox2* gene (Camus *et al.* 2017b).

*Mitochondrial physiology.*

Mitochondrial bioenergetics were characterized on fly permeabilized tissue through high resolution fluorespirometry, using two dedicated Oxygraphs-2k and DatLab v7.4 software (Oroboros Instruments, Innsbruck, Austria). Minor modifications were applied from existing protocols (Bettinazzi *et al.* 2019; Rodríguez *et al.* 2021; Rodríguez *et al.* 2023). Fly thoraces (2 thoraces per sample) were dissected on ice and subsequently permeabilized with the detergent saponin (80 µg·$mL^{-1}$) in BIOPS preservation solution (2.77 mM $CaK_2EGTA$, 7.23 mM $K_2EGTA$, 5.77 mM $Na_2ATP$, 6.56 mM $MgCl_2$, 20 mM taurine, 15 mM $Na_2$phosphocreatine, 20 mM imidazole, 0.5 mM dithiothreitol, and 50 mM K-MES, pH 7.1) for 30 minutes. After permeabilization, fly tissues were rinsed for 5 minutes in respiratory medium, carefully dried, weighted, and transferred into the O2k respiratory chambers set at 25°C. Chambers were preloaded with 2.1 mL MiR05 respiratory medium (110 mM D-sucrose, 60 mM lactobionic acid, 20 mM taurine, 20 mM HEPES, 10 mM $KH_2PO_4$, 3 mM $MgCl_2$, 0.5 mM EGTA, BSA 1 g·$L^{-1}$) (Gnaiger 2020). Before each experiment, the oxygen level was calibrated at air saturation, while the fluorescence signal with known concentration of $H_2O_2$ (0, 0.1 and 0.2 µM), in presence of 15 µM DTPA, 5 U·$mL^{-1}$ superoxide dismutase (SOD), 1 U·$mL^{-1}$ horseradish peroxidase (HRP), and 10 µM Amplex Ultra Red (AmR). Chambers where then sealed and both $O_2$ and $H_2O_2$ fluxes measured in parallel at 25°C, following a specific substrate-uncoupler-inhibitors-titration (SUIT) protocol. Mitochondrial respiration linked with the futile proton cycle in absence of ADP (Leak, L-state - state 4 or 2’) was assessed in presence of the complex I (CI)-specific substrates pyruvate (10 mM) and malate (2 mM) ($CI_L$). Supplementing ADP (5 mM) promoted mitochondrial respiration (OXPHOS, P-state or state 3) sustained by CI substrates ($CI_P$). Coupled respiration was then further fuelled by the stepwise addition of the substrates proline (10 mM), succinate (10 mM) and glycerophosphate (10 mM), respectively stimulating the additional activity of proline dehydrogenase (ProDH) ($CI{+}ProDH_P$), complex II (CII) ($CI{+}ProDH{+}CII_P$) and glycerophosphate dehydrogenase (GpDH) ($CI{+}ProDH{+}CII{+}GpDH_P$), the latter parameter representing the max coupled respiration achieved in this protocol. Sequential titration of the protonophore FCCP (0.125 µM each step) promoted the max uncoupled respiration (ETS, E-state or state 3u) ($CI{+}ProDH{+}CII{+}GpDH_E$). Uncoupled respiration in absence of CI and CII was then sequentially measured in presence of 0.5 µM rotenone (Rot - complex I inhibitor) ($ProDH{+}CII{+}GpDH_E$) and 5 mM malonate (Mal – complex II inhibitor) ($ProDH{+}GpDH_E$). Addition of 2.5 µM antimycin A (Ama - complex III inhibitor) resulted in the residual oxygen consumption (ROX). The standalone activity of cytochrome *c* oxidase (complex IV, CIV) was assessed in presence of 2 mM ascorbate and 0.5 mM TMPD, and corrected for the residual oxygen consumption with 20 mM sodium azide (Azd - complex IV inhibitor) ($CIV_E$). All respirometry data were corrected for the instrumental background measured in separate experiments. $H_2O_2$ fluxes were evaluated in parallel with respiratory rates, and parameters named accordingly (tables s1,s3).

Respirometry data were expressed as $O_2$ fluxes (pmol $O_2 \cdot s^{-1} \cdot mg^{-1}$), normalized for tissue mass, or as flux control ratios (FCR), normalized for the max coupled respiration (Gnaiger 2020). Comparison of thoraxes mass (mg $\cdot$ 2 thorases$^{-1}$), as well as raw flux data (pmol $O_2 \cdot s^{-1} \cdot mL^{-1}$) were also reported for completeness. Line specific mitochondrial phenotypes were also condensed by means of a principal component analysis which combined all the $O_2$ flux parameters (figure s3, table s1). The first principal component accounted for 59.9% of the variability and provides a proxy of the maximal respiratory capacity, with the max coupled and uncoupled respiration, and CIV activity heavily loading on it (figure s3A,B,C,E). The total variability explained by the second principal component was 31.6%, mostly reflecting the respiration rates sustained by CI- and CII-linked substrates, with parameters $CI_P$, $CI+ProDH_P$ and $CI+ProDH+CII_P$ mostly contributing to it (figure s3A,B,D,F). The change in respiration from a background ($Y_x$) to a stimulated ($Z_x$) state, produced by the addition of a specific substrate (x) was expressed by means of flux control factors (FCF). Formula: $x_{CF} = 1-(Yx \cdot Zx^{-1})$. Control factor parameters were obtained for the substrates ADP, proline, succinate, glycerophosphate (G3P) and rotenone. Finally, the CIV apparent excess capacity indicates the exceeding capacity of cytochrome *c* oxidase over the max coupled respiration. Formula: $j_{ExCIV} = (CIV_E \cdot CI+ProDH+CII+GpDH_P{}^{-1})-1$ (table s1). Reactive oxygen species rates were expressed as $H_2O_2$ fluxes, normalized for tissue mass (pmol $H_2O_2 \cdot s^{-1} \cdot mg^{-1}$), or as ratios, in function of the step-specific oxygen consumption $((H_2O_2 \cdot O_2{}^{-1})\%)$ ($n$ = 6 for each experimental group - tT, mT, tM, mM - females and males) (table s3).

*mtDNA copy number.*

Total DNA was extracted with the DNeasy Blood & Tissue Kit (Qiagen) from a pool of 3 flies per sample, and mtDNA abundance determined fluorometrically on a Mastercycler® RealPlex thermocycler (Eppendorf, DE), using the KAPA SYBR® FAST qPCR Master Mix Kit (KAPABIOSYSTEMS). The reaction (20 µL) consisted in 1X KAPA buffer 1X, 0.2 µM per primer and 10 ng template. Initial denaturation was set at 95°C for 3 min, followed by 40 cycles of 95°C for 5 s, 60°C for 10 s and 72°C for 10 s, and by a final extension at 72°C for 1 min. Copy number was determined in duplicate using two complementary set of primers. One designed to amplify *cox1* mitochondrial gene: cox1-F (TCCTGATATAGCATTCCCACG), cox1-R (CAACTGAAGCTCCACCATGA), and the other the nuclear gene *rosy*: rosy-F

(GGTGGTGAGCCTGTTCTTCAAG), rosy-R (ACTGGTGTGTGGAATGTCTCGG). For both genes, the cycle thresholds (CT) – i.e. the number of cycles reflecting the intersection between the amplification curve and the instrumental threshold line - were measured in duplicates. Replicates with more than 13% difference were discarded and a negative control was run in absence of sample. The mtDNA copy number relatively to the nuclear genome was determined by the formula $(2^{-\Delta CT})\cdot 2$, with ΔCT referring to the difference between the mitochondrial and the nuclear gene mean CT ($n$ = 9 and $n$ = 10 for each experimental group - tT, mT, tM, mM - females and males, respectively) (table s2) (Ballard *et al.* 2007).

*Locomotor activity.*

Fly locomotor activity was monitored using dedicated Drosophila Activity Monitors (DAM2, Trikinetics) (Chiu *et al.* 2010; Anderson *et al.* 2022). Each DAM2 tube (5 mm diameter) was prepared *a priori* by adding food (1:1 P:C diet) to one end, which also prevented fly desiccation. Female and male flies belonging to the four mitonuclear lines were then anesthetised ($CO_2$), added to each tube and let acclimate for half a day before starting the experiment. Tubes were then randomly placed into the monitors, where an infrared beam monitored the fly passages at the centre of tube. Fly activity was recorded for 48 hours at 25°C, 50% RH, under 12:12 light:dark cycles, and calculated as the number of counts (beam breaks) per minute. The activity was then condensed in 30 minutes activity, and further in timeframe-specific activity (i.e. dawn, day, dusk and night) ($n$ = 62-64 and $n$ = 63-64 for each experimental group - tT, mT, tM, mM - females and males, respectively). The data presented refers to the end of day 1 and day 2 of monitoring (table s4).

*Female fecundity, fertility and eggs to adults survival rate.*

Experimental females of each mitonuclear line (tT, mT, tM and mM) and standard males (LHm strain), all virgins, were collected 2 hours after eclosion, placed in separated vials at a concentration of 15 individuals per vial and let acclimate for three days (25°C, 50% RH, 1:1-P:C diet, 12:12 light:dark cycle). Vials containing four days old adults were then paired (15 females with 15 males) and let mate for 5 hours. Focal females were then placed in separated vials, supplemented with dry yeast *ad libitum*, and let lay eggs for a period of 14 hours. After that, females were discarded and eggs counted (fecundity - n eggs · female$^{-1}$). Vials with no eggs were discarded. Eggs were then let develop for 12 days at standard conditions and the adult offspring counted (fertility - n adults · female$^{-1}$). The eggs to adult survival rate was also calculated ((adults · eggs$^{-1}$) %) ($n$ = 56-60 for each experimental group - tT, mT, tM, mM - females) (table s5) (Tanaka & Yamazaki 1990; Camus *et al.* 2017a; Camus *et al.* 2020b).

*Male fertility (non-competitive).*

Experimental males of each mitonuclear strain (tT, mT, tM, mM) and standard females (LHm strain) were collected into separate vials at a concentration of 15 individuals per vial. After three days of acclimation, focal males of each strain were placed along with standard LHm females (15 focal males with 15 standard females for each pairing). All adults were virgins, reared at the same conditions (25°C, 50% RH, 1:1-P:C diet, 12:12 light:dark cycle) and of the same age (four days old). Flies were then allowed to interact for 5 hours. Females were then sorted in individual vials and let them lay eggs for 48 hours. After that time, they were transferred to new vials and let oviposit for additional 48 hours. Eggs were left develop for 12 days at standard conditions and the adult offspring in each vial counted (fertility - n adults · female$^{-1}$). Vials with no adults were discarded ($n$ = 37-39 for each experimental group - tT, mT, tM, mM - males) (table s6) (Camus *et al.* 2020a).

*Male fertility (competitive).*

Experimental males of each mitonuclear line (tT, mT, tM, mM), competitor males (LHm *bw-*) and standard females (LHm *bw-*), were separated into different vials at a concentration of 20 individuals per vial and let acclimate at standard conditions (25°C, 50% RH, 1:1-P:C diet, 12:12 light:dark cycle). All adults were virgins, reared at the same conditions and of the same age (5 days old). For each mitonuclear line, a trio of experimental males were placed together with a trio of LHm *bw-* competitor males and six LHm *bw-* females. Individuals were let interact for 24 hours, time during which focal and rival males competed for mating with females. Adults were then discarded, and eggs let develop for 12 days, period after which the progeny was counted and sorted based on the eye colour. Given that the competitor males and the standard females were both of LHm genetic background and homozygous for the recessive *bw−* eye-colour allele (brown eyes), whereas experimental adults are of the wild type (red eyes), the progeny with red eyes were assigned to the focal males (tT, mT, tM or mM), whereas flies with brown eyes were assigned to the competitor line. Male fertility was defined as the percentage of red-eyes adults of the total offspring yield (fertility (WT · adults$^{-1}$) %). Vials with no adults were discarded (*n* = 20 for each experimental group - tT, mT, tM, mM - males) (table s7) (Collet *et al.* 2016; Camus *et al.* 2017a).

*Development time and survival.*

Four days old flies of each mitonuclear line (tT, mT, tM, mM) were placed in separate oviposition chambers containing an agar–grape juice media with *ad libitum* live yeast, and let mate - lay eggs for 2 hours. Eggs were then collected and placed in separate vials at a concentration of 30 eggs per vial, up to a total of 20 vials for each mitonuclear genotype.

Three time a day and until all individuals developed, vials were screened for newly hatched adults. Adults were then removed and both development time and sex recorded (development time - hours) ($n$ = 206-280 and $n$ = 177-224 for each experimental group - tT, mT, tM, mM - females and males, respectively). Survival to adulthood was also measured as the number of successfully hatched adults over the starting number of eggs in the vial (survival – (adults · eggs$^{-1}$) %) ($n$ = 20 for each experimental group - tT, mT, tM, mM – females and males) (tables s8,s9) (Rodríguez *et al.* 2021).

*Heat challenge.*

Four days old flies of each sex and mitonuclear genotype were placed in separate 5 mL water-tight glass vials and immersed in a circulating water bath preheated at 39°C. The position of flies of each experimental group was randomized in each experiment. Heat tolerance was measured as the time (min) taken for each fly to enter a coma-like state (heat knock-down) (Hoffmann *et al.* 2002; Camus *et al.* 2017b). The experiment was performed over five trials within two different generations, and each trial consisted of a fully balanced replicate of the experimental ($n$ = 50 for each experimental group - tT, mT, tM, mM - females and males) (table s10).

*Cold challenge.*

Flies of each sex and mitonuclear genotype were placed in separate 1.5 mL tubes and immersed in an ice-slurry water bath prechilled at 0°C for 4 h, to place flies into a cold-induced coma. Tubes were then removed and laid out at 25°C. Recovery time was measured as the time (min) taken for each fly to regain consciousness after the coma (CCRT - chill-coma recovery time) (Camus *et al.* 2017b). For graphical purposes, cold tolerance was expressed as 120 minus CCRT. The experiment was performed over 6 trials within two different generations, and each trial consisted of a fully balanced replicate of the experimental ($n$ = 60 for each experimental group - tT, mT, tM, mM - females and males) (table s11).

*Data analysis.*

Graphical representation and data analysis were performed using the R software (R Core Team 2021) and several packages including 'ggplot2' (Wickham 2016), 'lme4' (Bates *et al.* 2015), 'lmerTest' (Kuznetsova *et al.* 2017), 'lsmeans' (Lenth 2016), 'multcomp' (Hothorn *et al.* 2008), 'car' (Fox & Weisberg 2019), 'FactoMineR' (Lê *et al.* 2008) and 'emmeans' (Lenth 2022). Shapiro and Levene's tests were respectively used to verify the normality and homoscedasticity of data. A linear mixed model was implemented for each parameter separately, considering the factors mitochondrial genome (*'mtDNA'*), nuclear background (*'nDNA'*), sex (*'sex'*) as categorical fixed effects, as well as their two-way (*'mtDNA:nDNA'*, *'mtDNA:sex'*, *'nDNA:sex'*) and three-way (*'mtDNA:nDNA:sex'*) interactions. The factors population (*'pop'* – i.e. different replicates of each population), generation (*'batch'*), fly age (*'day'*), trial (*'run'*) and vial (*'vial'*) were included as random effects. A generalized linear mixed model fitting the same fixed and random effects was implemented for parameters following either Poisson (female fecundity and fertility, male fertility) or binomial (egg-adult viability; WT·adults$^{-1}$) distribution. For egg-adult survival parameters (from either female fitness or development assays), the binomial model included flies that failed to develop (deaths) as a response variable. For male fitness (competitive) parameter, the binomial model included brown- and red-eyed adults as a response variable. The significance of the three factors and their possible interactions were determined through a Type III ANOVA, followed by *post hoc* multi comparison and Hommel adjustment for multiple testing. Starting from a full model, the best fitting model was determined through a step-wise model simplification by backward elimination of nonsignificant highest order effects. Statistical significance was set at $p \leq 0.05$. Data are graphically presented as mean ± standard error of the mean (s.e.m). Detailed summaries are provided in supplementary tables s1-s11. Datasets and R scripts for all analyses can be found on figshare online repository: doi:10.6084/m9.figshare.24162879.

### REFERENCES


1. Adrion, J.R., Hahn, M.W. & Cooper, B.S. (2015). Revisiting classic clines in Drosophila melanogaster in the age of genomics. *Trends Genet*, 31, 434-444.

2. Anderson, L., Camus, M.F., Monteith, K.M., Salminen, T.S. & Vale, P.F. (2022). Variation in mitochondrial DNA affects locomotor activity and sleep in Drosophila melanogaster. *Heredity*, 129, 225-232.

3. Ballard, J.W., Melvin, R.G., Miller, J.T. & Katewa, S.D. (2007). Sex differences in survival and mitochondrial bioenergetics during aging in Drosophila. *Aging cell*, 6, 699-708.

4. Bates, D., Mächler, M., Bolker, B. & Walker, S. (2015). Fitting Linear Mixed-Effects Models Using lme4. *Journal of Statistical Software*, 67, 1 - 48.

5. Bergland, A.O., Tobler, R., González, J., Schmidt, P. & Petrov, D. (2016). Secondary contact and local adaptation contribute to genome-wide patterns of clinal variation in Drosophila melanogaster. *Molecular Ecology*, 25, 1157-1174.

6. Bettinazzi, S., Rodríguez, E., Milani, L., Blier, P.U. & Breton, S. (2019). Metabolic remodelling associated with mtDNA: insights into the adaptive value of doubly uniparental inheritance of mitochondria. *Proceedings of the Royal Society B: Biological Sciences*, 286, 20182708.

7. Camus, M.F., Fowler, K., Piper Matthew, W.D. & Reuter, M. (2017a). Sex and genotype effects on nutrient-dependent fitness landscapes in *Drosophila melanogaster*. *Proceedings of the Royal Society B: Biological Sciences*, 284, 20172237.

8. Camus, M.F., Moore, J. & Reuter, M. (2020a). Nutritional geometry of mitochondrial genetic effects on male fertility. *Biology Letters*, 16, 20190891.

9. Camus, M.F., O'Leary, M., Reuter, M. & Lane, N. (2020b). Impact of mitonuclear interactions on life-history responses to diet. *Philos Trans R Soc Lond B Biol Sci*, 375, 20190416.

10. Camus, M.F., Wolff, J.N., Sgro, C.M. & Dowling, D.K. (2017b). Experimental Support That Natural Selection Has Shaped the Latitudinal Distribution of Mitochondrial Haplotypes in Australian *Drosophila melanogaster*. *Mol Biol Evol*, 34, 2600-2612.

11. Chakraborty, A., Sgrò, C.M. & Mirth, C.K. (2020). Does local adaptation along a latitudinal cline shape plastic responses to combined thermal and nutritional stress? *Evolution*, 74, 2073-2087.

12. Chiu, J.C., Low, K.H., Pike, D.H., Yildirim, E. & Edery, I. (2010). Assaying locomotor activity to study circadian rhythms and sleep parameters in Drosophila. *J Vis Exp*.

13. Clancy, D.J. (2008). Variation in mitochondrial genotype has substantial lifespan effects which may be modulated by nuclear background. *Aging cell*, 7, 795-804.

14. Collet, J.M., Fuentes, S., Hesketh, J., Hill, M.S., Innocenti, P., Morrow, E.H. *et al.* (2016). Rapid evolution of the intersexual genetic correlation for fitness in Drosophila melanogaster. *Evolution*, 70, 781-795.

15. Fox, J. & Weisberg, S. (2019). *An R Companion to Applied Regression*. Third edition edn. Sage, Thousand Oaks CA.

16. Gnaiger, E. (2020). *Mitochondrial Pathways and Respiratory Control An Introduction to OXPHOS Analysis, 5th Edition*. Bioenergetics Communications.

17. Hoffmann, A.A., Anderson, A. & Hallas, R. (2002). Opposing clines for high and low temperature resistance in Drosophila melanogaster. *Ecology Letters*, 5, 614-618.

18. Hoffmann, A.A. & Weeks, A.R. (2007). Climatic selection on genes and traits after a 100 year-old invasion: a critical look at the temperate-tropical clines in Drosophila melanogaster from eastern Australia. *Genetica*, 129, 133-147.

19. Hothorn, T., Bretz, F. & Westfall, P. (2008). Simultaneous inference in general parametric models. *Biom J*, 50, 346-363.

20. Kuznetsova, A., Brockhoff, P.B. & Christensen, R.H.B. (2017). lmerTest Package: Tests in Linear Mixed Effects Models. *Journal of Statistical Software*, 82, 1 - 26.

21. Lajbner, Z., Pnini, R., Camus, M.F., Miller, J. & Dowling, D.K. (2018). Experimental evidence that thermal selection shapes mitochondrial genome evolution. *Sci Rep*, 8, 9500.

22. Lê, S., Josse, J. & Husson, F. (2008). FactoMineR: An R Package for Multivariate Analysis. *Journal of Statistical Software*, 25, 1 - 18.

23. Lenth, R.V. (2016). Least-Squares Means: The R Package lsmeans. *Journal of Statistical Software*, 69, 1 - 33.

24. Lenth, R.V. (2022). _emmeans: Estimated Marginal Means, aka Least-Squares Means_. R package version 1.8.2.

25. R Core Team (2021). R: A language and environment for statistical computing. *R Foundation for Statistical Computing, Vienna, Austria*.

26. Rodríguez, E., Bettinazzi, S., Inwongwan, S., Camus, M.F. & Lane, N. (2023). Harmonising protocols to measure Drosophila respiratory function in mitochondrial preparations. *Bioenergetics Communications*, 2023.3.

27. Rodríguez, E., Grover Thomas, F., Camus, M.F. & Lane, N. (2021). Mitonuclear Interactions Produce Diverging Responses to Mild Stress in Drosophila Larvae. *Frontiers in genetics*, 12.

28. Sgro, C.M., Overgaard, J., Kristensen, T.N., Mitchel, K.A., Cockerell, F.E. & Hoffmann, A.A. (2010). A comprehensive assessment of geographic variation in heat tolerance

and hardening capacity in populations of *Drosophila melanogaster* from eastern Australia. *Journal of Evolutionary Biology*, 23, 2484-2493.

29. Tamura, K., Stecher, G. & Kumar, S. (2021). MEGA11: Molecular Evolutionary Genetics Analysis Version 11. *Molecular Biology and Evolution*, 38, 3022-3027.

30. Tanaka, T. & Yamazaki, T. (1990). Fitness and its components in *Drosophila melanogaster*. *The Japanese Journal of Genetics*, 65, 417-426.

31. Wickham, H. (2016). *ggplot2: Elegant Graphics for Data Analysis.* Springer-Verlag New York.

32. Zhu, C.-T., Ingelmo, P. & Rand, D.M. (2014). G×G×E for Lifespan in *Drosophila*: Mitochondrial, Nuclear, and Dietary Interactions that Modify Longevity. *PLOS Genetics*, 10, e1004354.

### SUPPLEMENTARY FIGURES AND TABLES

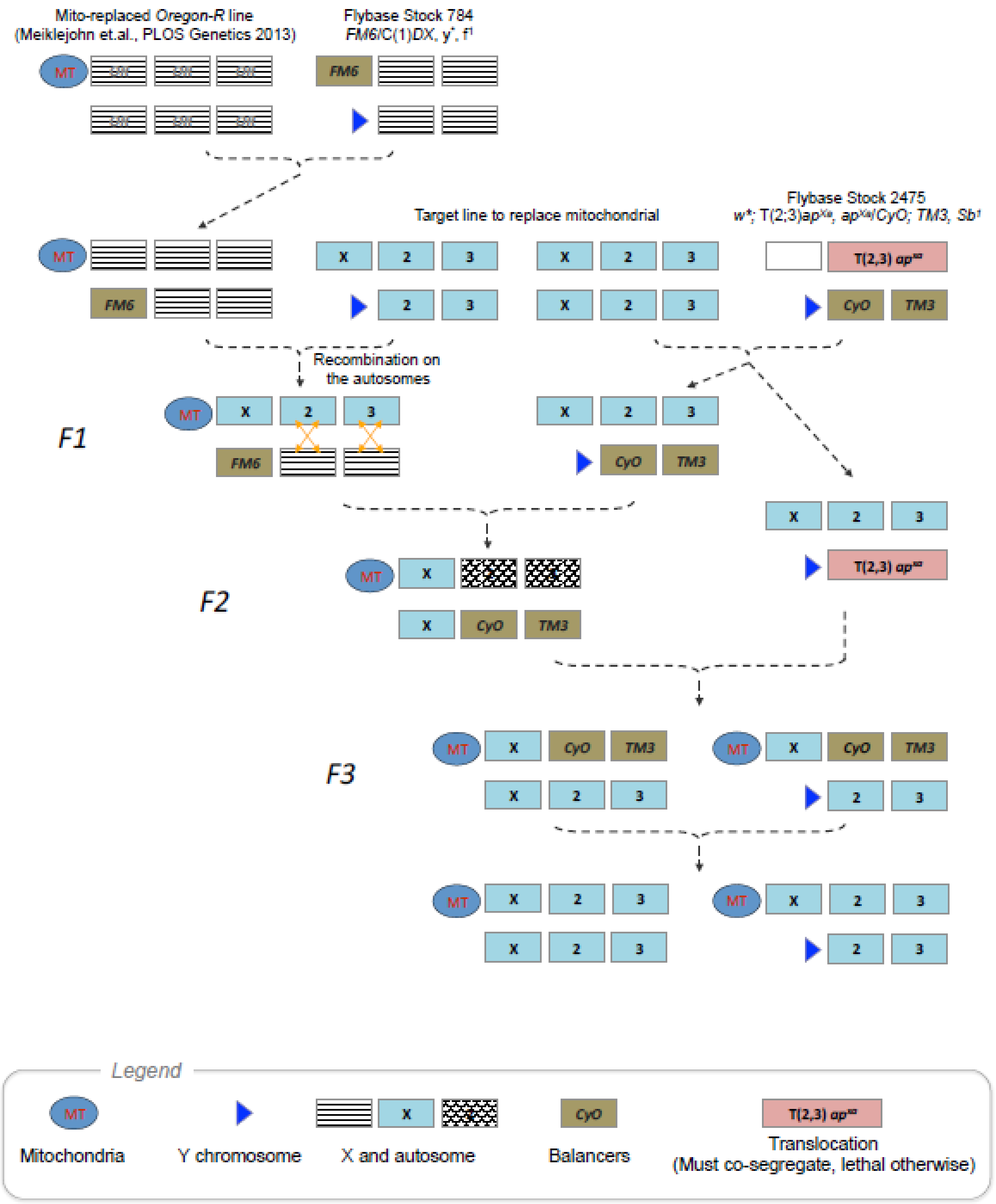


**Figure s1. Schematics for mitochondrial replacement by balancer substitution.**

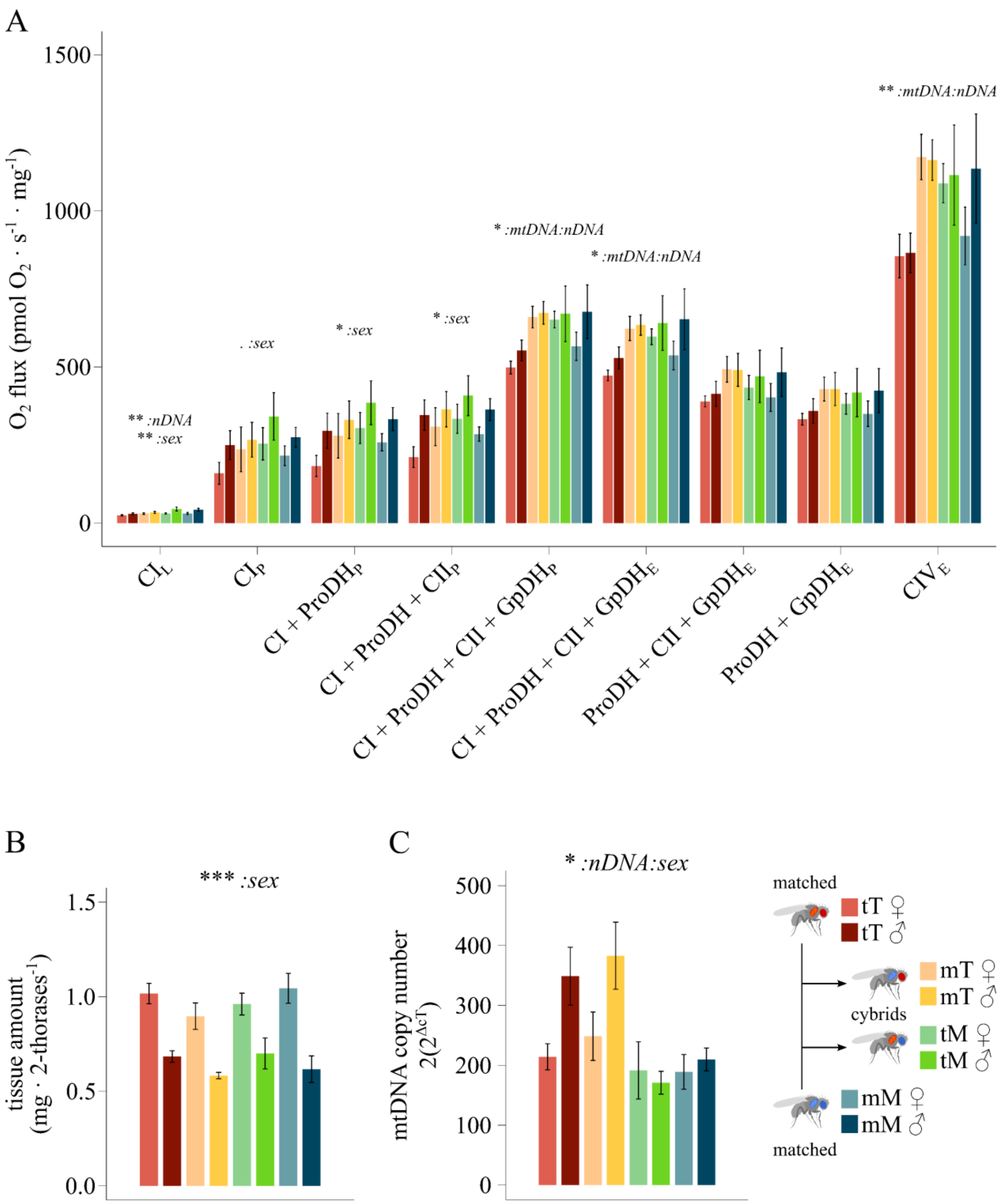


**Figure s2. Mitochondrial respiratory fluxes, tissue wet weight and mtDNA copy number. (A)** Mitochondrial respiratory fluxes in permeabilized fly thoraces normalized for tissue mass (pmol $O_2$ · $s^{-1}$ · $mg^{-1}$) (*n*=6). **(B)** Tissue amount (mg) per sample (2 thoraces) (*n*=6). **(C)** Mitochondrial DNA copy number ($2(2^{\Delta cT})$) (*n*=9,10). *Parameters: 'CI$_L$'*: CI-sustained leak respiration; *'CI$_P$'*: CI-linked coupled respiration; *'CI+ProDH$_P$'*: CI+ProDH-linked coupled respiration; '*CI+ProDH+CII*$_P$': CI+ProDH+CII-linked coupled respiration; '*CI+ProDH+CII+GpDH*$_P$': max coupled respiration with CI+ProDH+CII+GpDH-linked substrates; '*CI+ProDH+CII+GpDH*$_E$': max uncoupled respiration with CI+ProDH+CII+GpDH-linked substrates; '*ProDH+CII+GpDH*$_E$': ProDH+CII+GpDH-linked uncoupled respiration; '*ProDH+GpDH*$_E$': ProDH+GpDH-linked uncoupled respiration; '*CIV*$_E$': cytochrome *c* oxidase standalone capacity. Statistical analyses: linear mixed model; Fixed effects: *'mtDNA'*, *'nDNA'* and *'sex'*, plus their interactions. Significance was determined by means of a type III ANOVA. Data shown as mean ± sem. *$p \leq 0.05$; **$p \leq 0.01$; ***$p \leq 0.001$. A detailed summary is reported in supplementary tables s1,s2.

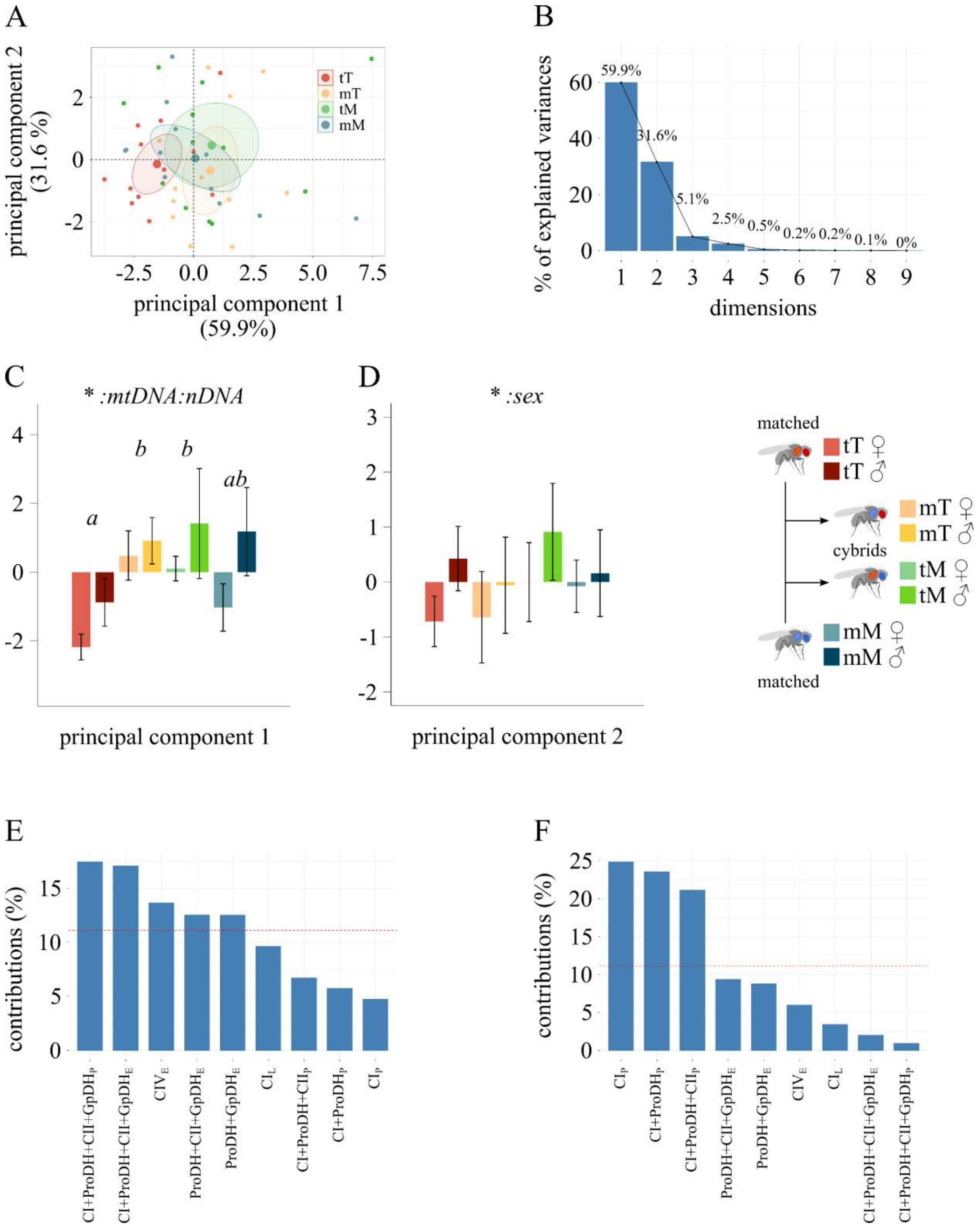


**Figure s3. Principal component analysis.** Principal component analysis (PCA) based on all respiratory fluxes normalized for tissue mass (pmol $O_2 \cdot s^{-1} \cdot mg^{-1}$) (figure s2A). **(A)** PCA scatter plot with 95% confidence interval ellipses. Colours refer to the different mitonuclear lines, comprising both female and males. **(B)** Percentage of explained variance for each principal component. **(C)** First principal component (PC1) (*n*=6). **(D)** Second principal component (PC2) (*n*=6). **(E)** Contribution of variables to PC1. **(F)** Contribution of variables to PC2. *Parameters:* '$CI_L$': CI-sustained leak respiration; '$CI_P$': CI-linked coupled respiration; '$CI+ProDH_P$': CI+ProDH-linked coupled respiration; '$CI+ProDH+CII_P$': CI+ProDH+CII-linked coupled respiration; '$CI+ProDH+CII+GpDH_P$': max coupled respiration with CI+ProDH+CII+GpDH-linked substrates; '$CI+ProDH+CII+GpDH_E$': max uncoupled respiration with CI+ProDH+CII+GpDH-linked substrates; '$ProDH+CII+GpDH_E$': ProDH+CII+GpDH-linked uncoupled respiration; '$ProDH+GpDH_E$': ProDH+GpDH-linked uncoupled respiration; '$CIV_E$': cytochrome *c* oxidase standalone capacity. Statistical analyses: linear mixed model; Fixed effects: *'mtDNA'*, *'nDNA'* and *'sex'*, plus their interactions. Significance was determined by means of a type III ANOVA. Letters indicate statistical difference following a *post-hoc* multi comparison test, with *p*-values adjusted using Hommel's correction. Data shown as mean ± sem. $*p \leq 0.05$; $**p \leq 0.01$; $***p \leq 0.001$. A detailed summary is reported in supplementary table s1.

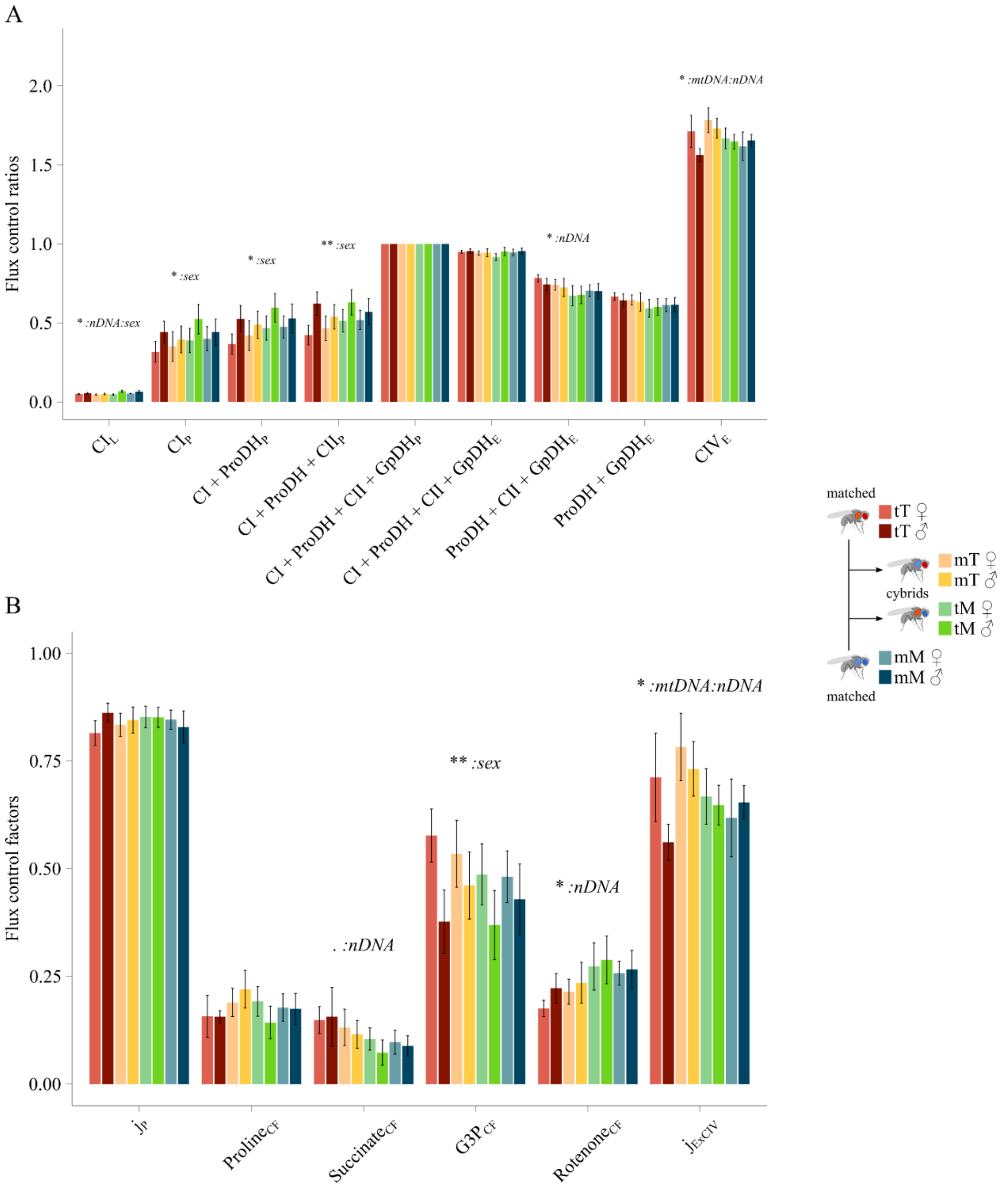


**Figure s4. Mitochondrial phenotype. (A)** Flux control ratios (FCR) expressing respirometry fluxes normalized for the max coupled respiration (CI+ProDH+CII+GpDH$_P$) ($n$=6). **(B)** Flux control factors (FCF), showing the change in respiration linked with the addition of a specific substrate ($n$=6). *Parameters: 'CI$_L$'*: CI-sustained leak respiration; *'CI$_P$'*: CI-linked coupled respiration; *'CI+ProDH$_P$'*: CI+ProDH-linked coupled respiration;

*'CI+ProDH+CII$_P$'*: CI+ProDH+CII-linked coupled respiration; *'CI+ProDH+CII+GpDH$_P$'*: max coupled respiration with CI+ProDH+CII+GpDH-linked substrates; *'CI+ProDH+CII+GpDH$_E$'*: max uncoupled respiration with CI+ProDH+CII+GpDH-linked substrates; *'ProDH+CII+GpDH$_E$'*: ProDH+CII+GpDH-linked uncoupled respiration; *'ProDH+GpDH$_E$'*: ProDH+GpDH-linked uncoupled respiration; *'CIV$_E$'*: cytochrome *c* oxidase standalone capacity; *'$j_P$'*: coupling efficiency; *'Proline$_{CF}$'*: proline control factor; *'Succinate$_{CF}$'*: succinate control factor; *'G3P$_{CF}$'*: glycerophosphate control factor; *'$j_{ExCIV}$'*: CIV excess capacity. Statistical analyses: linear mixed model; Fixed effects: *'mtDNA'*, *'nDNA'* and *'sex'*, plus their interactions. Significance was determined by means of a type III ANOVA. Data shown as mean ± sem. *$p \leq 0.05$; **$p \leq 0.01$; ***$p \leq 0.001$. A detailed summary is reported in supplementary table s1.

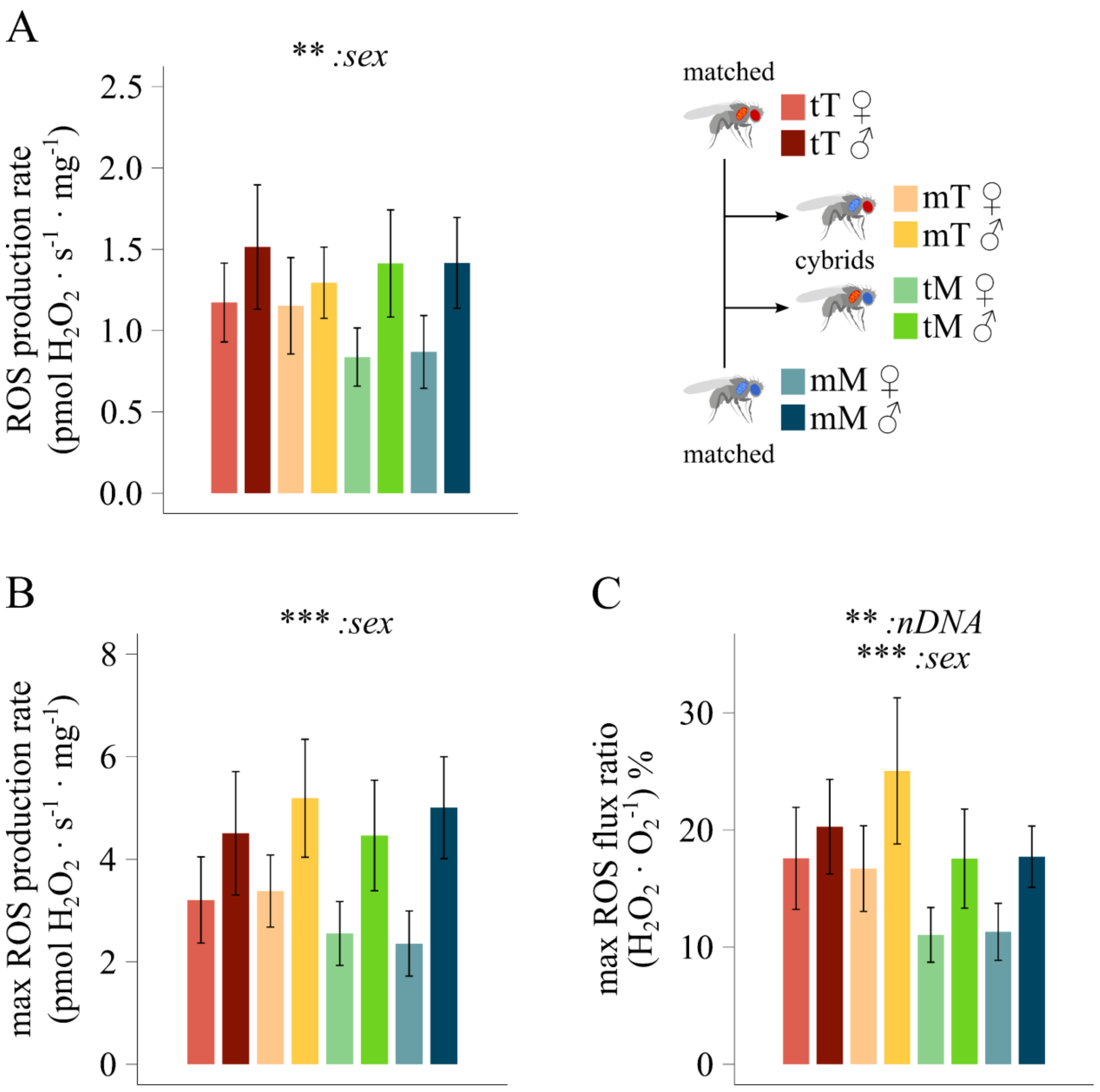


**Figure s5. Reactive oxygen species metabolism. (A)** Hydrogen peroxide production rate during max coupled respiration (pmol $H_2O_2 \cdot s^{-1} \cdot mg^{-1}$) (*n*=6). **(B)** Hydrogen peroxide maximal production rate following the complete inhibition of the electron transport system (pmol $H_2O_2 \cdot s^{-1} \cdot mg^{-1}$) (*n*=6). **(C)** Hydrogen peroxide production over oxygen consumption following the complete inhibition of the electron transport system (($H_2O_2 \cdot O_2^{-1}$) %) (*n*=6). Statistical analyses: linear mixed model; Fixed effects: *'mtDNA'*, *'nDNA'* and *'sex'*, plus their interactions. Significance was determined by means of a type III ANOVA. Data shown as mean ± sem. *$p \leq 0.05$; **$p \leq 0.01$; ***$p \leq 0.001$. A detailed summary is reported in supplementary table s3.

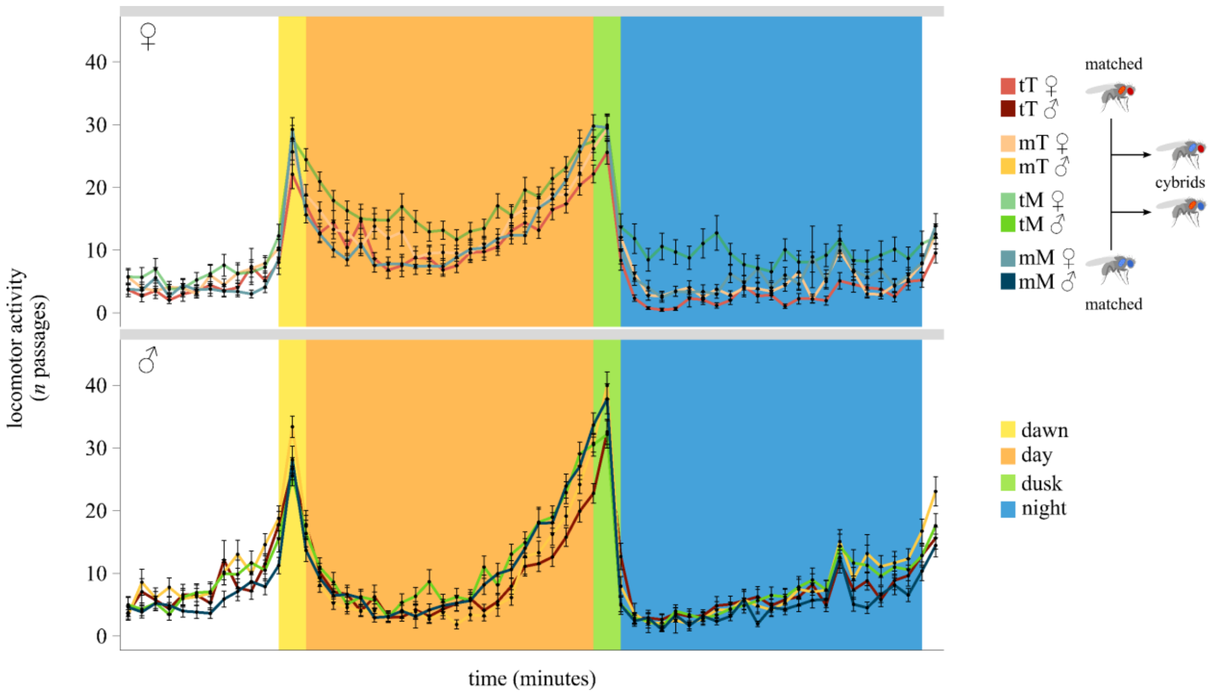


**Figure s6.** Locomotor activity (*n* passages) throughout one day (*n*=62-64). Each datapoint refers to a 30 minutes activity. Data are shown as mean ± sem. Coloured areas define different timeframes, respectively dawn, day, dusk and night. A detailed summary is reported in supplementary table s4.

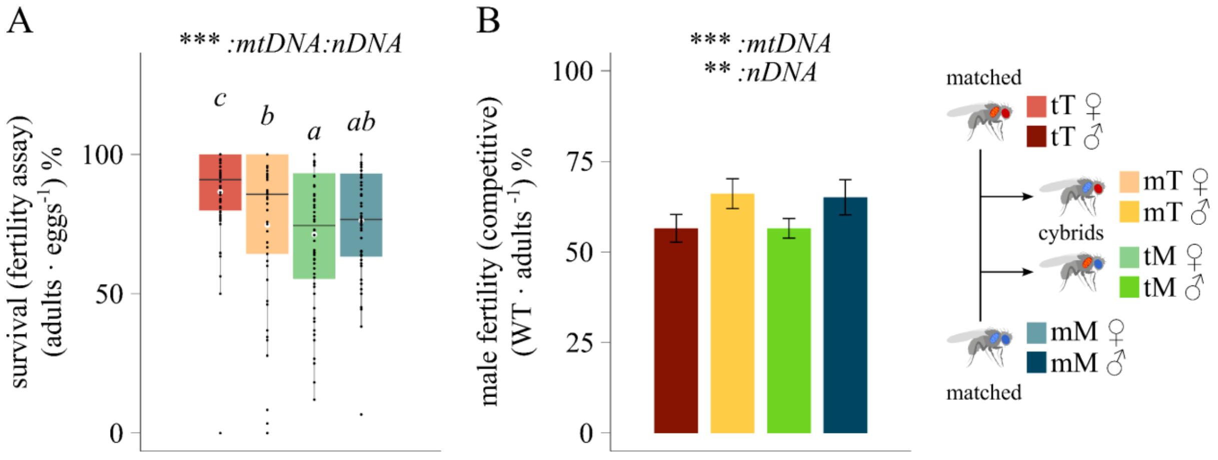


**Figure s7. Reproductive fitness. (A)** Female reproductive success expressed as egg-adult survival ((adults · eggs$^{-1}$) %) ($n$=56-60). **(B)** Male competitive reproductive success expressed as the percentage of wild-type (WT – red-eyes) offspring produced over the total number of adults. Standard LH$_m$ females mated with competing focal males and LHm males ((fertility - WT adults · tot adults$^{-1}$)%). Statistical analyses: generalized linear mixed model; Fixed effects: *'mtDNA'*, *'nDNA'* and *'sex'*, plus their interactions. Significance was determined by means of a type III ANOVA. Letters indicate statistical difference following a *post-hoc* multi comparison test, with $p$-values adjusted using Hommel's correction. Data shown as mean ± sem. *$p \leq 0.05$; **$p \leq 0.01$; ***$p \leq 0.001$. A detailed summary is reported in supplementary tables s5, s7.

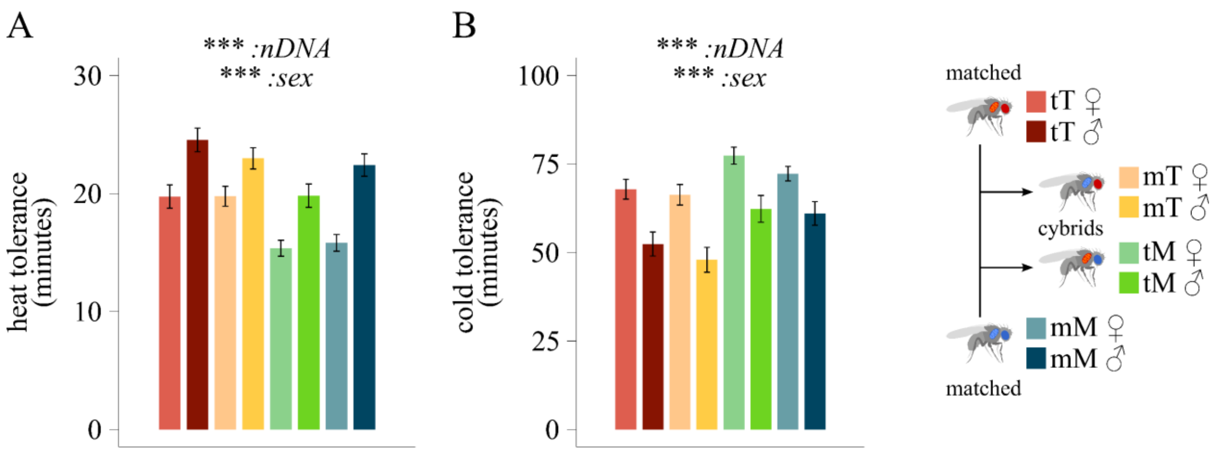


**Figure s8. Thermal tolerance. (A)** Heat shock tolerance measured as the time (min) taken for each fly to enter a coma-like state following 39°C heat shock ($n$=50). **(B)** Cold shock tolerance was expressed as 120 minus the time (min) taken for each fly to regain consciousness after chill-induced coma ($n$=60). Statistical analyses: linear mixed model; Fixed effects: *'mtDNA'*, *'nDNA'* and *'sex'*, plus their interactions. Significance was determined by means of a type III ANOVA. Data shown as mean ± sem. *$p \leq 0.05$; **$p \leq 0.01$; ***$p \leq 0.001$. A detailed summary is reported in supplementary tables s10-11.
